# Early resident NK cell response to local HIV infection in lymphoid tissue

**DOI:** 10.1101/2025.02.17.638403

**Authors:** David Perea, Alba Gonzalez, Nerea Sanchez-Gaona, Felix Pumarola, Nuria Ortiz, Ines Llano, Juan Lorente, Vicenç Falcó, Meritxell Genescà, Maria J. Buzon

## Abstract

Natural killer (NK) cells are critical mediators of antiviral immunity, yet their role in lymphoid tissues—key reservoirs of HIV persistence—remains poorly defined. Here, we uncover a distinct cytotoxic signature of tonsil-resident NK cells essential for targeting HIV-infected CD4⁺ T cells. Using a human tonsillar model of HIV infection and extracellular matrix-based functional assays, we identify a subset of NK cells co-expressing CD69, CD49a, CD103, and the adaptive marker NKG2C as potent effectors against autologous HIV-infected tissue CD4⁺ T cells. Both CD16⁺ and CD16⁻ NK subsets exhibited cytotoxic antiviral activity. However, HIV infection induced profound functional alterations in these NK cells, including dysregulated expression of *CD9, TRAF2, ITGA1*, suggesting disrupted activation, signaling, and tissue residency. Functional assays corroborated a significant impairment in the cytotoxic capacity, indicating HIV-driven NK cell dysfunction. Intriguingly, a subset of immature CD16⁻CD69⁺ NK cells underwent functional reprogramming, transitioning into a migration-competent, and metabolically primed state, driven by the upregulation of *NUMAI, LCP1, SLC38A1, MT-ND2, NUMA1, MYH9, and CD44*. Functional assays confirmed this transition, revealing changes in the expression of immune checkpoint receptors and a gain of cytotoxic function in these reprogrammed cells. These findings advance our understanding of NK cell biology in HIV infection and highlight novel avenues for NK-cell-based therapeutic strategies.

## INTRODUCTION

HIV primarily replicates in CD4^+^ T lymphocytes and causes AIDS if untreated. Presently, while antiretroviral therapy (ART) can reduce the viral load to undetectable levels, HIV remains an incurable condition^1^. During acute infection, HIV rapidly disseminates to secondary lymphoid tissues, where reservoirs are quickly established^2^. Even with ART, lymphoid tissue and the gastrointestinal tract act as key sanctuaries for HIV persistence, posing a major obstacle to a cure^3,4^.

Natural killer (NK) cells are innate lymphocytes (ILCs) that efficiently kill tumor and infected cells without prior antigen sensitization^5^; thus, the recruitment of functionally mature NK cells may efficiently control HIV^3^. In African green monkeys (AGMs), an animal model of SIV control, viral infection induces the expansion of terminally differentiated NK cells in secondary lymphoid tissues, which mediate a strong control of the infection^6^. The recruitment of cytotoxic NK cells has been linked to increased IL-15 production, specifically localized in B cell follicles of AGMs^7^. Importantly, mucosal NKp44^+^ ILCs have also demonstrated beneficial roles in controlling SIV infection^8,9^ and are found expanded in tonsils during chronic SIV infection^10^. In humans, elevated NK cell levels in lymph nodes have been observed during the first weeks of HIV infection^11,12^, contrasting with their scarce presence one year later due to impaired NK cell migration^3^. The slow recruitment of cytotoxic NK cells may facilitate the establishment of latently infected cells in lymphoid tissues, as HIV can reach lymph nodes within few days after infection^2^. Moreover, infected lymphoid tissue undergoes rapid structural remodeling^13,14^ and loss of functional integrity^15,16^, which are essential for mounting an effective immune response. Therefore, understanding the function of ILCs, more specifically NK cells, already residing within these tissues is crucial, as they may act as early innate immune responders to HIV infection, potentially limiting viral infection and the establishment of viral persistence.

Extensive research has been conducted on NK cells circulating in peripheral blood, where cytokines activate less differentiated CD56^+^CD16^−^ NK cells prompting them to quickly respond and eliminate abnormal cells^17^. Generally, their functional response is activated by surface receptors that transduce activating and inhibitory signals. Among these receptors, there are Killer Cell Ig-like Receptors (KIRs), Natural Cytotoxicity Receptors (NCRs; NKp30, NKp44, NKp46), C-type-Lectin Receptors (CLRs; NKG2A, NKG2C, NKG2D) as well as immune checkpoints (ICs; PD-1, TIGIT, KLRG1)^18–21^. Of special interest is the CD16 molecule, which links innate and adaptive immunities to selectively kill infected or abnormal cells by antibody-dependent cell-mediated cytotoxicity (ADCC)^17^. Moreover, CD16^+^ NK cells are highly cytotoxic and have been shown to play a critical role in immune surveillance, patrolling the bloodstream and tissues to identify and target aberrant cells^22^.

Circulating CD56^+^ NK cells can also home to tissues, giving rise to tissue-resident NK cells (trNK cells)^23–25^. Most tissues contain trNK cells, which express the markers involved in cell retention, such as CD69, CD49a, and CD103^23^. In general, trNK cells are typically less cytotoxic than blood NK cells^26^, and seem to produce higher levels of cytokines such as IFN-γ^27,28^. Animal models have shown that trNK cells proliferate locally in response to acute and chronic viral infections^29^, and can directly regulate the magnitude of T-cell specific immune response^30^. Notably, the expression of various resident markers is associated with distinct functional outcomes, as illustrated in the healthy lung^31^ and liver^32^. For example, CD56^+^CD16^−^CD49a^+^ NK cells produce elevated levels of proinflammatory cytokines but demonstrate poor degranulation^31,33^. Furthermore, recent studies indicate that the tissue site can also dictate the potential functions of trNK subsets. In a seminal study, Dogra *et al*. detailed the organ distribution of classic CD56^bright^CD16^−^ (proinflammatory) and CD56^dim^CD16^+^ (cytotoxic) NK subsets^23^, revealing tissue-specific divergences, with CD56^bright^CD16^−^ NK cells predominating in lymph nodes, tonsil, and the gut^23,34^. Others have further described tonsil-resident NK cells similar to circulating CD56^bright^ NK cells, mainly lacking perforin, KIRs, NKp30, and CD16, but partially expressing NKp44 and NKp46^34–37^. Moreover, trNK cells may exhibit organ-specific functions unique to different tissues. For instance, in the uterus, they contribute to placental vascular remodeling, fetal growth, and the establishment of pregnancy memory^38,39^. Although NK cells are primarily innate immune effectors, adaptive or memory-like NK cells have been observed to expand selectively during certain viral infections, including HIV^40,41^ ^42^. The most widely recognized marker of memory NK cells is NKG2C, which facilitates the recognition of specific viral antigens^42^. Additionally, the KIR repertoire is closely associated with adaptive NK cells, playing a crucial role in the education of NK cells prior to the acquisition of NKG2C^43^. With the exception of CD49a^+^ NK cells ^29,33,44^, this memory-like activity has been predominantly identified in circulating NK cells. Despite these significant advances in understanding NK cell functions, the role of different NK cell subsets during HIV infection, as well as the modulation of these cells by HIV in tissues, remains a pivotal yet unresolved question.

Here, using an *ex vivo* explant model of HIV-infected human tonsillar tissue, we demonstrate how the expression of tissue-resident memory markers and immune checkpoints influences the function of tonsillar ILCs cells during HIV infection. We identified distinct CD56^+^ cell populations with functional properties targeting both conventional K562 cells—the gold standard for assessing NK cell functionality in cell culture—and HIV-infected cells. Our findings reveal HIV-driven transcriptional changes that influence their effector functions. Additionally, our findings underscore the importance of assessing NK cell activity in more physiologically relevant models that preserve the tissue microstructure. This work provides crucial insights into the immune landscape of HIV infection in lymphoid tissues and highlights potential avenues for enhancing innate cell-mediated antiviral responses in therapeutic strategies.

## RESULTS

### Phenotypic characterization of tonsillar ILCs

We first performed a phenotypic characterization of NK cells present in tonsils. Tissue was obtained from 68 donors (**Supplementary Table S1)**. Non-B cell lymphocytes represented a median of 40.8% out of the CD45^+^CD14^−^CD19^−^ cells **(Fig. 1A, left)**. NK cells were identified by gating for CD45^+^CD3^−^CD14^−^CD19^−^CD56^+^ cells, which were further classified into CD16^+^ and CD16^−^ subsets **(Supplementary Fig. S1A-B)**. In general, NK cells within tonsils were infrequent, accounting for a median of approximately 0.7% of CD45^+^CD14^−^CD19^−^ lymphocytes **(Fig. 1A, middle)**, as previously reported by Dogra *et al*.^23^. Notably, the majority of these tonsillar CD56^+^ NK cells were CD16^−^ (median 73.1%) **(Fig. 1A, right)**. We then analyzed the coexpression of memory-like markers including KIRs^43^ and NKG2C^33^, and the residency markers CD69, CD49a, and CD103 in the two major NK subsets. We found that CD16^+^ NK cells displayed a higher prevalence of markers associated with tissue residency, specifically CD49a and CD103 **(Fig. 1B)**, as well as memory-like markers **(Fig. 1C)**. It is noteworthy that NK cells expressing CD69 were abundant in both the CD16^+^ and CD16^−^ subsets, with a notably higher frequency in the CD16^−^ subset (79.4%; *p*=<0.001) **(Fig. 1B)**. Moreover, CD49a^+^ (CD103^+/−^) NK cells exhibited a notable enrichment in the expression of memory-like markers, especially the CD16^+^ NK subset **(Fig. 1D, S1C)**.

**Figure 1.**
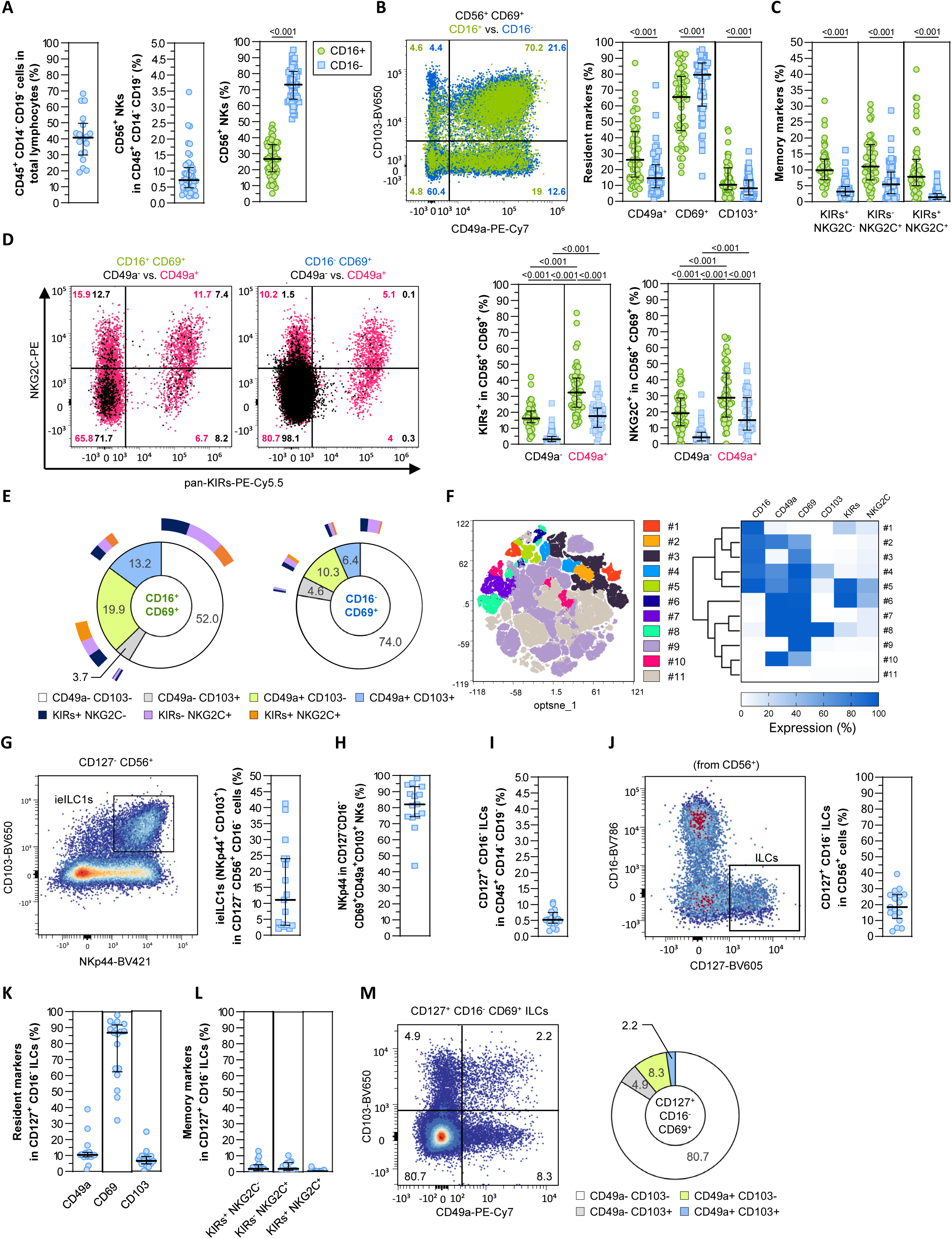
Tissue-resident memory phenotype of tonsillar NK cells. Expression of CD56, CD16, NKp44, resident (CD69, CD49a, and CD103) and memory (KIRs and NKG2C) markers was measured by flow cytometry in different tonsillar tissue specimens. **(A)** Frequencies of CD45^+^CD14^−^CD19^−^ lymphocytes (left), total CD56^+^ NK cells (middle) and classic CD16^+/−^ NK cell subsets (right). **(B)** Frequency of expression of resident markers within CD56^+^CD16^+/−^ subsets. **(C)** Frequency of expression of memory markers within CD56^+^CD16^+/−^ subsets. **(D)** Frequency of expression of memory markers within CD49a^+^ and CD49a^-^ NK cells. **(E)** Coexpression frequencies of resident and memory markers within CD56^+^CD16^+/−^ NK cell subsets. **(F)** Optimized-t-Distributed Stochastic Neighbor Embeddings (opt-SNEs) plot showing the abundance of tissue-resident memory NK cell clusters (#) within the total pool of CD56^+^ NK cells in tonsils at baseline. The heatmap shows the frequency of expression of different tissue-resident memory markers per cluster. **(G)** Frequency of ieILC1s (NKp44^+^CD103^+^ within CD127^−^CD56^+^CD16^−^ cells). **(H)** Frequency of NKp44 expression within CD127^−^CD56^+^CD16^−^CD69^+^CD49a^+^CD103^+^ NK cells. **(I-J)** Frequencies of CD127^+^CD16^−^ ILCs within **(I)** CD45^+^CD14^−^CD19^−^ lymphocytes and **(J)** CD56^+^ cells. **(K)** Frequency of resident CD127^+^CD16^−^ ILCs. **(L)** Frequency of memory CD127^+^CD16^−^ ILCs. **(M)** Coexpression frequencies of resident markers within CD127^+^CD16^−^ ILCs. For (A), (B), (C), (D), (G), (H), (I), (J), (K), and (L), median with interquartile range is represented. Statistical comparisons were performed using the Wilcoxon matched-pairs signed rank test. Significant comparisons *p*<0.05. Green: CD16^+^; blue: CD16^−^.

In the process of infiltrating into tissues, NK cells first upregulate CD69 expression, which interferes with sphingosine-1-phosphate receptor-1 (S1PR1) and effectively block lymphocyte egress^45,46^. Subsequently, the tissue microenvironment influences the expression of additional residency markers, including CD49a (collagen I-IV binding)^47^ and CD103 (E-cadherin binding)^48,49^, in response to local inflammation and cytokine cues from neighboring tissue cells^37,50,51^. Thus, we analyzed the co-expression of these residency markers in tonsillar tissue, since this might provide important localization and functional information. We observed that the majority of the NK cells in both CD16^+^ and CD16^−^ NK subsets were identified as CD69^+^CD49a^−^CD103^−^ (SP, for single-positive CD69), with a median of 52.0% for CD16^+^ and 74.0% for CD16^−^ **(Fig. 1E)**. A secondary, but still substantial, population of CD16^+^CD69^+^CD49a^+^CD103^−^ NK cells (DP, for double-positive CD69 and CD49a), was also observed, comprising approximately 20% of all CD16^+^CD69^+^ NK cells **(Fig. 1E)**. The majority of CD49a and CD103 coexpression was observed in CD69⁺ NK cells. Specifically, 13.2% of CD16⁺ NK cells and 6.4% of CD16⁻ NK cells expressed these markers (**Fig. 1E**). Finally, to gain further insights into the heterogeneity of tonsillar NK cells, we conducted a dimensionality reduction analysis (opt-SNE), which revealed the presence of 11 distinct cell clusters based on resident and memory-like markers **(Fig. 1F)**. In concordance with our previous results, clusters #3 and #9 collectively represented the majority of tissue-resident NK cells within the tonsil, exhibiting a phenotype characterized by the single expression of CD69. In contrast, clusters #4 and #8 corresponded to NK cells with triple resident marker expression (TP, for triple-positive CD69, CD49a, and CD103), while cluster #5 represented a unique subset of CD16^+^CD69^+^CD49a^+^ resident NK cells exhibiting high KIRs and NKG2C expression, and cluster #6 corresponded to its CD16^−^ counterpart.

We next evaluated the coexpression of CD103 and NKp44 **(Fig. 1H)**, confirming the existence of the previously described intraepithelial ILC1s (ieILC1s)^52^. These ieILC1s represented a median of 82% of all CD127^−^CD56^+^ CD16^−^ TP cells **(Fig. 1G)**. Based on these findings, we will now refer to the CD16^−^ TP cells as ieILC1s. Other ILCs were present within the tonsillar samples. Total CD127^+^CD16^−^ ILCs accounted for a median of 0.5% of CD45^+^CD14^−^CD19^−^ lymphocytes **(Fig. 1I)**, and within CD56^+^ cells, ILCs represented a median of 18.4% **(Fig. 1J)**. Importantly, resident **(Fig. 1K)** and memory **(Fig. 1L)** receptors were found to be rare on these ILCs, except for the abundant expression of CD69 observed (Fig. 1M).

In summary, while we identified CD56^+^CD16^−^CD69^+^CD49a^−^CD103^−^ (CD16^−^ SP) NK tissue subpopulation as predominant in tonsils, CD16^+^CD69^+^CD49a^+^ subset exhibited a significant expression of KIRs and NKG2C. This likely indicates a more mature NK cell phenotype, consistent with the well-established role of the tonsils as secondary lymphoid organs, where NK cells undergo differentiation^53^.

### Functionality of cytotoxic ILCs present in tonsillar tissue

We first determined the functional basal status of tonsillar NK cells based on the expression of the degranulation marker CD107a and the proinflammatory cytokine IFN-γ^54,55^. Additionally, their maximum functional degranulation and proinflammatory potentials were assessed as the frequency of CD107a^+^IFN-γ^+/−^ NK cells upon coculturing with the MHC class I-devoid K562 cell line, the gold standard for measuring NK cell functionality in culture. Initially, we observed that CD16^+^ NK cells exhibited low expression of CD107a **(Fig. 2A)**. Conversely, CD16^−^ NK cells, in general, showed elevated basal CD107a expression (median of 4.7% vs 45.6% of CD107a^+^IFN-γ^−^ cells in the CD16^+^ and CD16^−^ NK subsets, respectively, *p*<0.001) **(Fig. 2A)**. Next, we focused on the CD49a^+^ population, as previous studies have identified this subset as one of the most functionally significant across various tissues^33,56^. The analysis revealed that in the absence of target cells, both CD16^−^ and CD16^+^ NK cells expressing CD49a displayed notably low levels of CD107a expression **(Fig. 2B, left)**. However, upon coculture with K562 target cells, degranulation was significantly upregulated, with the CD16^−^ CD49a^+^ subset exhibiting higher CD107a expression compared to the CD16^+^ fraction (median of 10.8% and 17.4% CD107a^+^IFN-γ^−^ cells in the CD16^+^ and CD16^−^ NK subsets, respectively) (**Fig. 2B, left**). In contrast, at baseline, the CD49a^−^ fraction within the CD16^−^ NK cells was predominantly CD107a^+^ (median of 55.6% CD107a^+^IFN-γ^−^ cells), and no significant upregulation of CD107a expression was observed following cell targeting (**Fig. 2B, right**). This suggests that this CD49a^−^ subset represents a functionally activated NK cell population in the tonsils, albeit with limited capacity for further stimulation. Additionally, we evaluated the ability of CD49a^+^ cells to produce IFN-γ following coculture with target cells. We found that both CD16^+^ and CD16^−^ subsets were capable of producing significant amounts of IFN-γ, with higher levels observed in the CD16^−^ fraction (**Fig. 2C, left**). In contrast, CD49a^−^ cells barely produce IFN-γ, either with or without cell stimulation (**Fig. 2C, right**). Overall, we found that CD49a^+^ NK cells, particularly within the CD16^−^ subset, exhibit high degranulation and IFN-γ production capacity, while CD49a^−^ cells are already expressing high levels of CD107a at baseline and show limited further response to stimulation.

**Figure 2.**
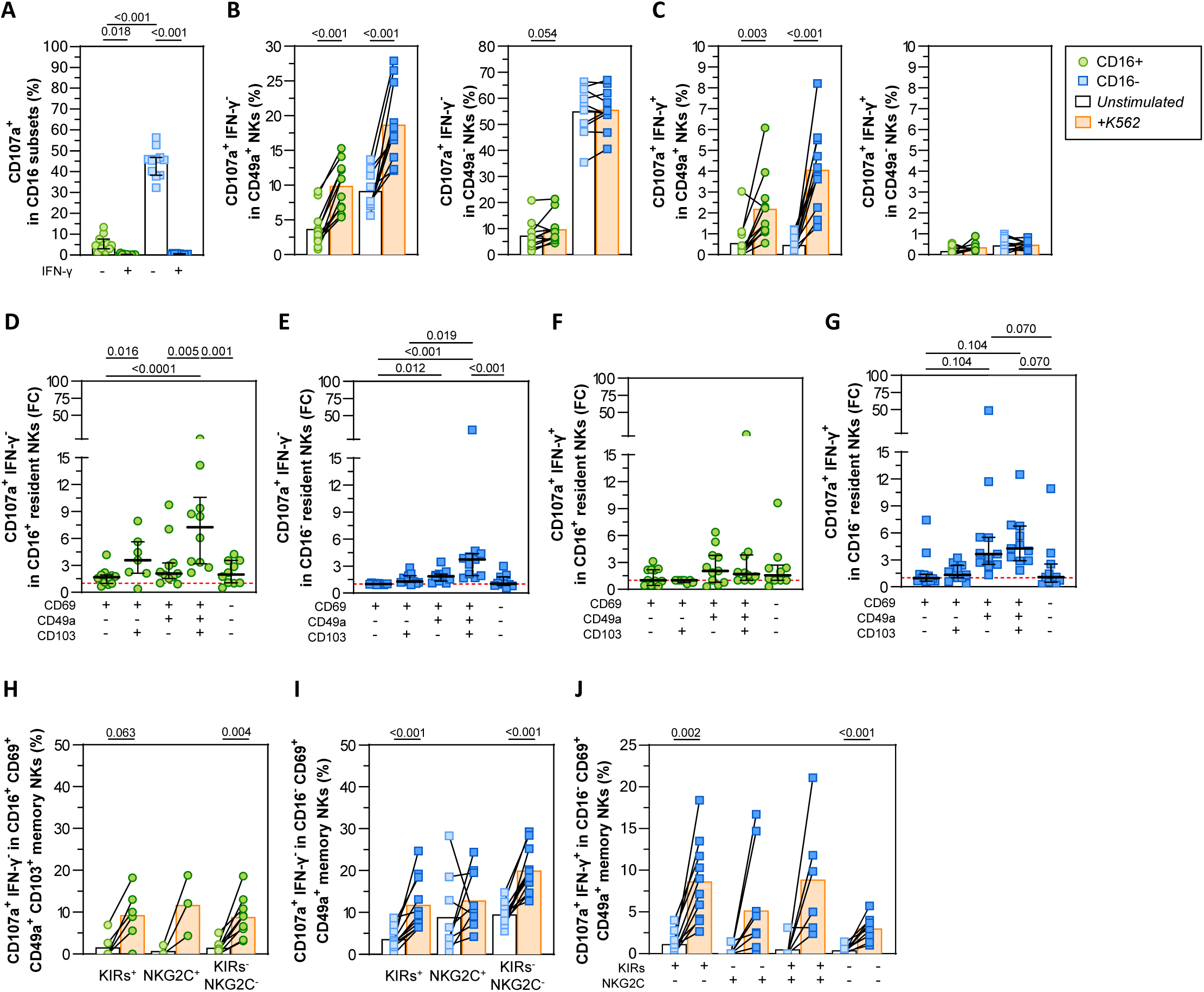
Functionality of tonsil-resident memory NK cells. Expression of the degranulation marker CD107a and intracellular IFN-γ measured by flow cytometry in different tonsillar tissue specimens in the steady state and upon stimulation with K562 cells. **(A)** Basal functional activation of classic CD16^+/−^ NK cell subsets. **(B)** Cytotoxic degranulation (% of CD107a^+^IFN-γ^−^ cells) by resident CD16^+/−^ CD49a^+^ (left) and CD49a^−^ (right) NK cells. **(C)** Cytokine production (% of CD107a^+^IFN-γ^+^ cells) by resident CD16^+/−^ CD49a^+^ (left) and CD49a^−^ (right) NK cells. **(D-E)** Cytotoxic degranulation (fold-change of % of CD107a^+^IFN-γ^−^ cells) by resident **(D)** CD16^+^ and **(E)** CD16^−^ NK cells. **(F-G)** Cytokine production (fold-change of % CD107a^+^IFN-γ^+^ cells) by resident **(F)** CD16^+^ and **(G)** CD16^−^ NK cells. **(H-I)** Cytotoxic degranulation (% of CD107a^+^IFN-γ^−^ cells) by **(H)** memory CD16^+^CD69^+^CD49a^+^CD103^+^ NK cells and **(I)** memory CD16^−^CD49a^+^ NK cells. **(J)** Cytokine production (% of CD107a^+^IFN-γ^+^ cells) by memory CD16^−^CD69^+^CD49a^+^ NK cells. Statistical comparisons were performed using the Wilcoxon matched-pairs signed rank test, Friedman test, and Skillings-Mack test when appropriate. Significant comparisons *p*<0.05. Red dotted line in fold-change graphs represents FC=1 (no change). Green circles: CD16^+^; blue squares: CD16^−^.

Next, we aimed to determine the contribution of CD103 in the functionality of CD49^+^ cells. In the CD16^+^ fraction, we found that CD107a production following cell encounters was predominantly achieved by the CD103^+^ population, which exhibited the highest degranulation capacity (FC = 7.3) upon K562 coculture (**Fig. 2D**). In contrast, CD16^−^ counterparts showed little increase in CD107a production (**Fig. 2E**) when expressing CD103. Regarding IFN-γ production, CD103 expression in the CD16^+^ fraction did not enhance its production following cell encounters (**Fig. 2F**). In the CD16^−^ fraction, however, CD49^+^ cells— both with and without CD103 expression—were identified as the most potent producers of IFN-γ (**Fig. 2G**), suggesting that CD103 may play a limited role in this specific cell function. A detailed analysis of all NK resident populations is provided in **Supplementary Fig. S2A-B**. Overall, we found that CD103 identifies cells with high cytotoxic potential within the CD16^+^ NK cell population, while no such role was observed in the CD16^−^ fraction.

Next, we assessed whether KIRs and NKG2C expression could identify functional signatures in the most active subsets identified above (all resident-memory NK populations are shown in **Supplementary Fig. 2C-F**). We found that the proportion of cells prone to degranulation alone (measured as CD107a^+^IFN-γ^−^) was not significantly altered in either the CD16^+^CD69^+^CD49a^+^CD103^+^ (**Fig. 2H**) or the CD16^−^CD69^+^CD49a^+^ cells upon expression of KIRs or NKG2C (**Fig. 2I**). However, the analysis revealed that in CD16^−^CD69^+^CD49a^+^ cells, coexpression of KIRs identified the most responsive subpopulations prone to both degranulation and IFN-γ secretion (**Fig. 2J**). These results suggest that KIRs expression serves as a clear indicator of cells with a more polyfunctional profile. This partially aligns with the work of Brownlie *et al*.^33^ in the lung, who observed that CD16^−^CD49a^+^KIRs^+^NKG2C^+^ NK cells are more responsive responded better than CD49a^−^ NK cells to K562 target cells.

In summary, we found that tonsillar CD16^−^CD69^+^CD49a^−^ NK cells, the most abundant population, have the highest basal CD107a production. However, upon K562 targeting, CD16^+^ CD69^+^CD49a^+^CD103^+^ cells were the most prone to degranulation and likely the most cytotoxic, while CD16^−^CD69^+^CD49a^+^KIR^+^ cells were identified as the more polyfunctional and cytokine-producing subset.

### Immune checkpoint signatures in tonsil-resident cytotoxic ILCs

Recognizing the distinct functionalities of cytotoxic and proinflammatory ILCs across CD16^+^ and CD16^−^ subsets, as well as resident memory populations, we analyzed the expression of various immune checkpoints (ICs) that may influence their functional fate (**Fig.S3A**). Inhibitory NKG2A is of particular interest, as HIV viral proteins can alter the expression of its ligand, HLA-E^57,58^. Additionally, LAIR-1 and KLRG1 are important inhibitory receptors for tissue-specific proteins, with LAIR-1 binding to collagens^59^ and KLRG1 targeting epithelial E-cadherin^60^. Moreover, we measured the expression of two other widely studied ICs, TIGIT and PD-1, which are known to be upregulated in memory NK cells in chronic HIV infection^61^.

Overall, tonsillar CD56^+^ cells exhibited high expression of NKG2A, LAIR-1, and TIGIT, while PD-1 and KLRG1 levels were relatively low **(Fig 3A)**. The CD16^+^ subset displayed elevated expression of all ICs except LAIR-1, which was more pronounced in the CD16^−^ population **(Fig. 3A)**. The division of CD56^+^ cells into different resident subpopulations revealed distinct IC expression patterns. Interestingly, CD16^+^ resident NK cells had elevated levels of NKG2A and TIGIT **(Fig. 3B)**, whereas CD16^−^ cells showed a progressive increase in NKG2A and TIGIT from the CD16^−^ SP to TP resident populations **(Fig. 3C)**. In contrast, LAIR-1 expression followed an inverse pattern in both CD16^+^ and CD16^−^ subsets **(Fig. 3B,C)**. PD-1 and KLRG1 expression remained low overall, though KLRG1 was more prominent in CD16^+^ SP NK cells **(Fig. 3B,C)**. We next focused on the expression of the ICs in the more functional populations identified earlier: the CD16^+^CD69^+^CD49a^+^CD103^+^ subset and the CD16^−^CD69^+^CD49a^+^KIR^+^ subset. For the CD16^+^CD69^+^CD49a^+^CD103^+^ tissue-resident TP NK cells, which produce high levels of CD107a upon cell stimulation, we observed that cells had low expression of LAIR and high expression of both NKG2A and TIGIT (**Fig. 3B**). While the low LAIR expression might support cytotoxic functions, the high levels of NKG2A and TIGIT may indicate mechanisms are in place to limit overactivation. For the CD16⁻CD69⁺CD49a⁺KIRs⁺ subset, which produces the highest levels of IFN-γ upon stimulation, we observed intermediate expression of all three ICs, NKG2A, LAIR, and TIGIT (**Fig. 3D**), suggesting that the co-expression of these inhibitory receptors on IFN-γ-producing cells may reflect a state of activation that is simultaneously regulated to prevent excessive immune responses. Expression of all ICs in the rest of resident memory NK cell populations is shown in **Supplementary Fig. S3A-C**.

**Figure 3.**
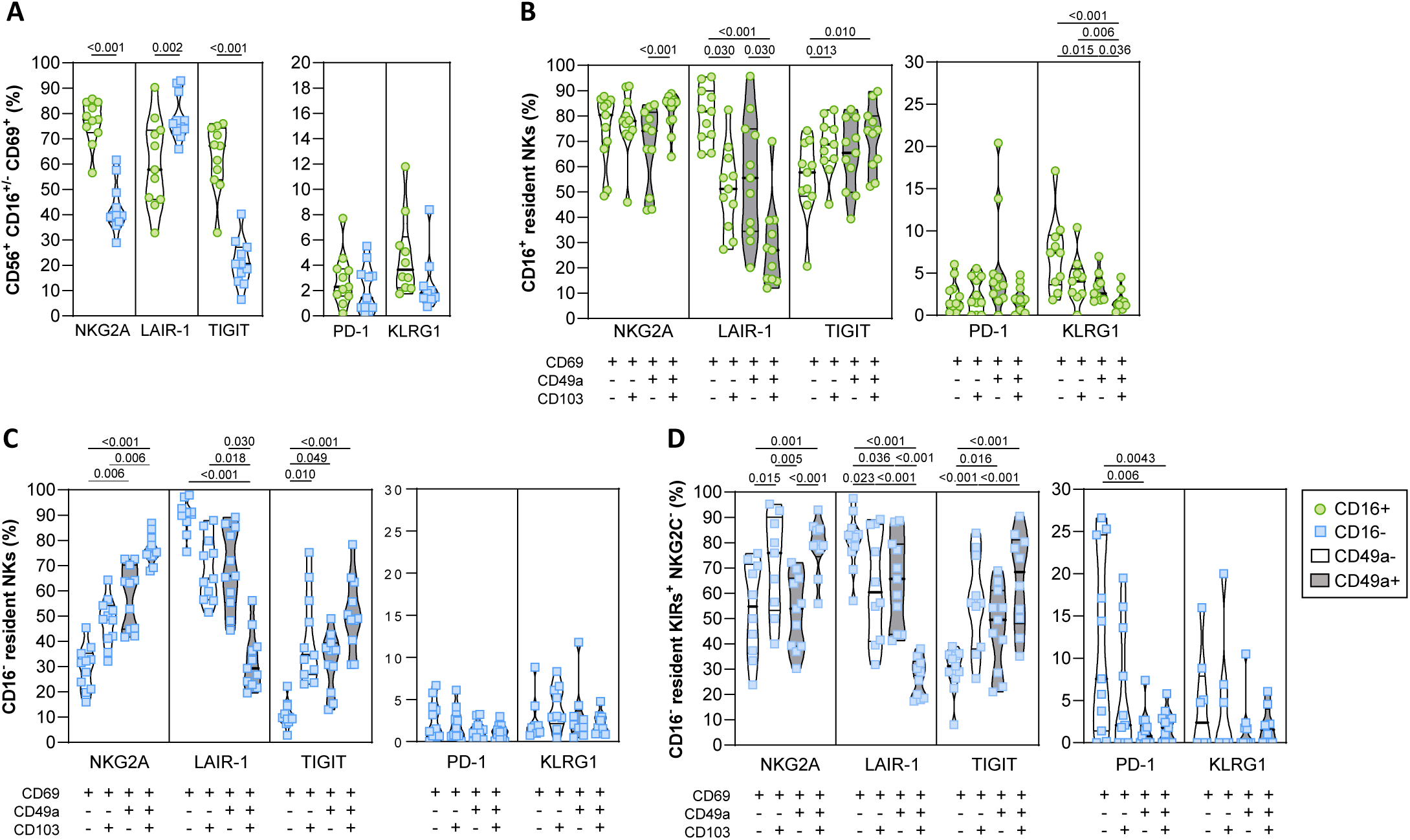
Immune checkpoint expression patterns in tonsil-resident NK cells. Expression levels of the inhibitory immune checkpoints NKG2A, LAIR-1, TIGIT, PD-1, and KLRG1 were analyzed by flow cytometry in tonsillar tissue samples. **(A)** Frequencies of immune checkpoint expression within CD56^+^CD16^+/−^ subsets. **(B-C)** Frequencies of immune checkpoint expression within resident CD16^+^ **(B)** and CD16^−^ **(C)** NK cells. **(D)** Frequencies of immune checkpoint expression within memory CD16^−^KIR^+^ NK cells. Statistical comparisons were performed using the Wilcoxon matched-pairs signed rank test or the Friedman’s test, as appropriate. Significant comparisons *p*<0.05. Green circles: CD16^+^; blue squares: CD16^−^.

In conclusion, the functional behavior of tonsil-resident NK cells is closely linked to the expression of immune checkpoint receptors, likely revealing different developmental and functional states.

### Extracellular matrix components modify the function of tonsil-resident NK Cells

The gold standard for evaluating the functional potential of NK cells involves coculturing them with K562 target cells in suspension and measuring functional surrogate markers following stimulation^62^. While this method is most likely well-suitable for the study of circulating NK cells, it is also usually used for tissue NK cells^33,44,63^ despite the growing evidence indicating that the extracellular matrix components in tissues might have an impact on NK cell functionality^64^. To account for this, we conducted functional assays with K562 cells incorporating 2D extracellular matrix components into the experimental setup (**Fig. 4A**). For the analyses, we focused on the populations that express CD49a, a collagen receptor with high affinity for basal lamina collagen IV (Col IV) and lower affinity for collagen IV (Col I). Thus, this receptor enables tissue scanning and facilitates interactions within the extracellular matrix^65^. We found that when NK cells were embedded in either Col I or Col IV, those expressing the CD49a marker exhibited a significantly higher capacity to produce CD107a alone, or both CD107a and IFN-γ. This was observed in both CD16^+^ and CD16^−^ subsets (**Fig. 4B**). In contrast, the same populations lacking CD49a expression did not show an increase in the production of the functional markers after coculturing with target cells in either collagen type (**Fig. 4C**). This suggests that CD49a plays a crucial role in not only NK cell activation but also in promoting effective immune responses within tissue matrices. Next, we focused on the more functional populations identified earlier: the CD16^+^CD69^+^CD49a^+^CD103^+^ subset, which primarily degranulates in response to target cells, and the CD16^−^CD69^+^CD49a^+^KIR^+^ subset, which produces IFN-γ upon target cell engagement. Notably, both populations express CD49a. We observed that CD16^+^CD69^+^CD49a^+^CD103^+^ cells exhibited increased degranulation in most experiments conducted with both collagens (**Fig. 4D**). Similarly, CD16^−^CD69^+^CD49a^+^KIR^+^ NK cells showed a notable increase in IFN-γ production when cocultured with K562 cells on collagen-coated wells (**Fig. 4E**). In contrast, the same subsets lacking CD49a did not respond to K562 cells (**Fig 3F-G**). Altogether, our findings indicate that extracellular matrix components alter NK functionality; particularly the capacity of CD49a^+^ NK cells to degranulate and produce IFN-γ in 2D collagen environments.

**Figure 4.**
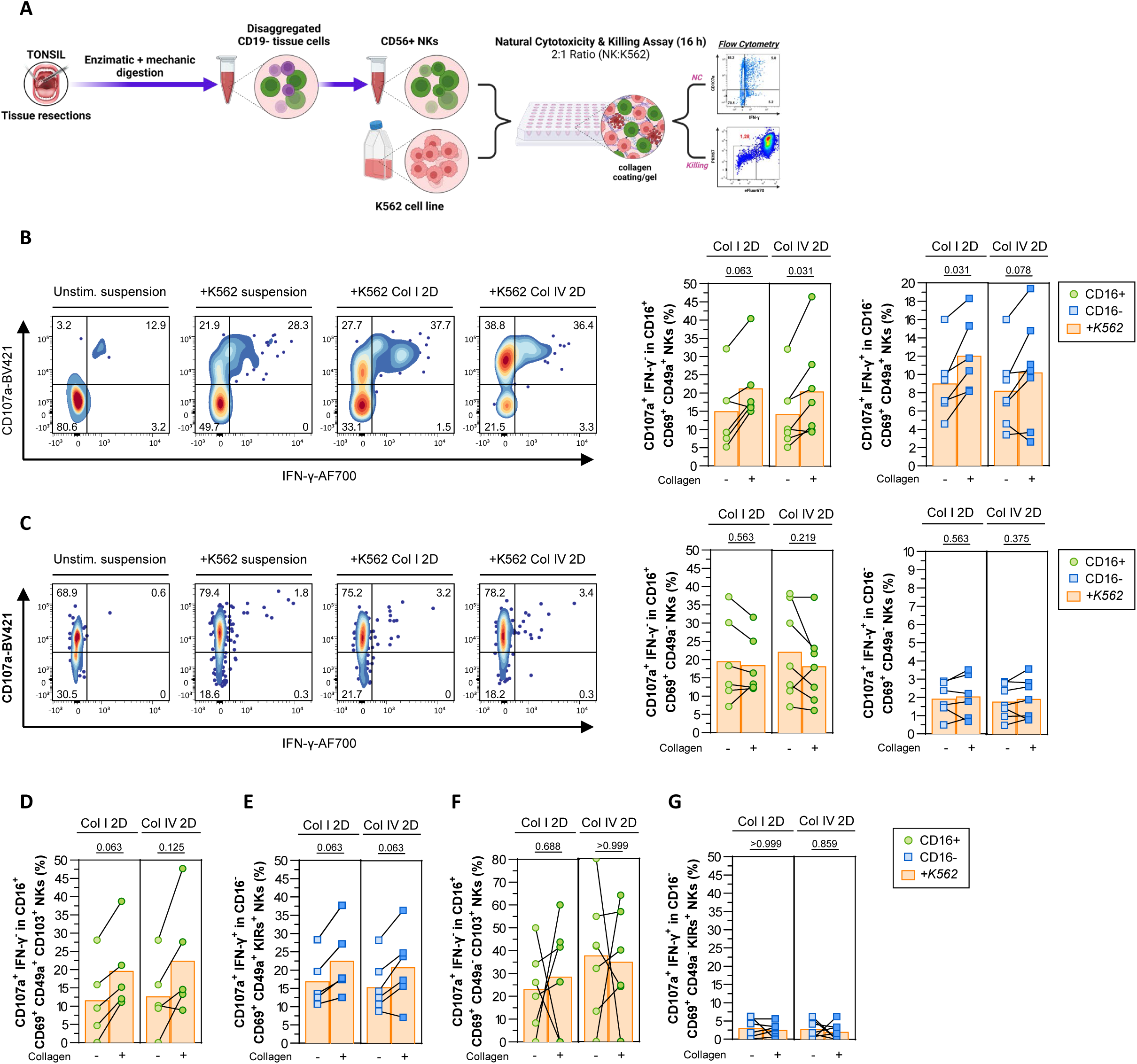
Role of the tissue microstructure in the function of resident NK cells. The effects of extracellular matrix components (collagens) were assessed in a functional coculture assay of tonsillar NK cells against K562 cell targets (2:1 ratio) overnight (16 hours). **(A)** Diagram illustrating the functional coculture assay of tonsil-resident NK cells in contact with extracellular matrix components. **(B)** Representative flow cytometry plots of functional markers expression within CD16^−^CD69^+^CD49a^+^ NK cells. Cytotoxic degranulation (% of CD107a^+^IFN-γ^−^) by CD16^+^CD69^+^CD49a^+^ NK cells and cytokine production (% of CD107a^+^IFN-γ^+^) by CD16^−^CD69^+^CD49a^+^ NK cells when cocultured with K562 target cells in suspension, on collagen I-or IV-coated surfaces (10 µg/cm^2^) at a cell density of 70,000 NK vs. 35,000 K562 cells per cm^2^. **(C)** Representative flow cytometry plots of functional markers expression within CD16^−^CD69^+^CD49a^−^ NK cells. Cytotoxic degranulation (% of CD107a^+^IFN-γ^−^) by CD16^+^CD69^+^CD49a^−^ NK cells and cytokine production (% of CD107a^+^IFN-γ^+^) by CD16^−^CD69^+^CD49a^−^ NK cells when cocultured with K562 target cells in suspension, on collagen I-or IV-coated surfaces (10 µg/cm^2^) at a cell density of 70,000 NK vs. 35,000 K562 cells per cm^2^. **(D)** Cytotoxic degranulation (% of CD107a^+^IFN-γ^−^) by CD16^+^CD69^+^CD49a^+^CD103^+^. **(E)** Cytokine production (% of CD107a^+^IFN-γ^+^) by CD16^−^CD69^+^CD49a^+^KIRs^+^. **(F)** Cytotoxic degranulation (% of CD107a^+^IFN-γ^−^) by CD16^+^CD69^+^CD49a^+^CD103^+^. **(G)** Cytokine production (% of CD107a^+^IFN-γ^+^) CD16^−^CD69^+^CD49a^−^KIRs^+^. NK cells when cocultured with K562 target cells in suspension, on collagen I-or IV-coated surfaces (10 µg/cm^2^) at a cell density of 70,000 NK vs. 35,000 K562 cells per cm^2^. Statistical comparisons were performed using the Wilcoxon matched-pairs signed rank test or the Friedman’s test, as appropriate. Significant comparisons *p*<0.05.

### Specific tonsil-resident memory NK cells are associated with enhanced killing of HIV-infected tonsillar CD4^+^ T cells

We next focused on identifying ILC populations involved in restraining HIV infection in the tonsillar model. To this end, we first examined the relationship between the frequencies of tonsil-resident NK cells and the levels of HIV infection. The median infection rate, represented by the percentage of CD3^+^CD8^−^ T cells expressing the viral antigen p24, was 1.0% (**Fig. S4A**), consistent with other reports^66^. We found that none of the resident markers in the CD16^+^ fraction were solely associated with improved control of HIV infection in tissue explants (**Fig. 5A**). However, a more detailed analysis revealed that higher frequencies of TP NK cells expressing NKG2C, and to lesser extend KIRs, within the CD16⁺ subset correlated with lower infection levels (**Fig. 5B**). Notably, the CD16⁺ TP population was identified as one of the most functional against K562 cells, as evidenced by CD107a production, with KIR and NKG2C expression not being critical for this feature. In contrast, against HIV, the expression of NKG2C appears to strongly correlate with lower infection levels. Moreover, the expression of ICs negatively impacted these CD16^+^ TP NK cells. Specifically, higher levels of p24 in the explants were associated with the expression of PD-1 **(Fig. 5C)** and lo a lesser extend KLRG1 **(Fig. 5D)** on these NK cells. For CD16⁻ NK cells, those expressing CD103 (CD49a⁺/⁻) exhibited beneficial effects (**Fig. 5E**), likely including ieILC1s, as most CD16⁻CD103⁺ cells express NKp44 (**Fig. 1G**). Importantly, CD16⁻CD103⁺ cells co-expressing CD49a (TP) correlated with lower infection levels when they also expressed NKG2C (**Fig. 5F**), similar to the CD16⁺ population. However, CD16⁻CD103⁺ cells lacking CD49a expression demonstrated the highest correlation against HIV when expressing KIRs (**Fig. 5F**). Furthermore, PD-1 expression on CD16⁻ TP NKG2C⁺ NK cells correlated with higher p24 levels (**Fig. 5G**), while KLRG1 expression on CD16⁻CD49a⁻CD103⁺ cells was also negatively associated with infection control in explants (**Fig. 5H**). Although the expression of most ICs on resident NK cells generally correlated with poorer HIV control, LAIR-1 showed an inverse pattern—its expression on less differentiated CD16⁺ (**Fig. S4B-C**) and CD16⁻ (**Fig. S4D-E**) NK cells was associated with lower HIV levels. Overall, we found that different ILC populations were associated with lower levels of HIV infection. Among all cells (CD16⁺/⁻), those expressing all three resident markers along with NKG2C showed this association. Interestingly, these cells exhibited the lowest levels of LAIR-1 while maintaining high levels of NKG2A and TIGIT (**Fig. 3D, S3B**), likely indicating a heightened readiness to respond.

**Figure 5.**
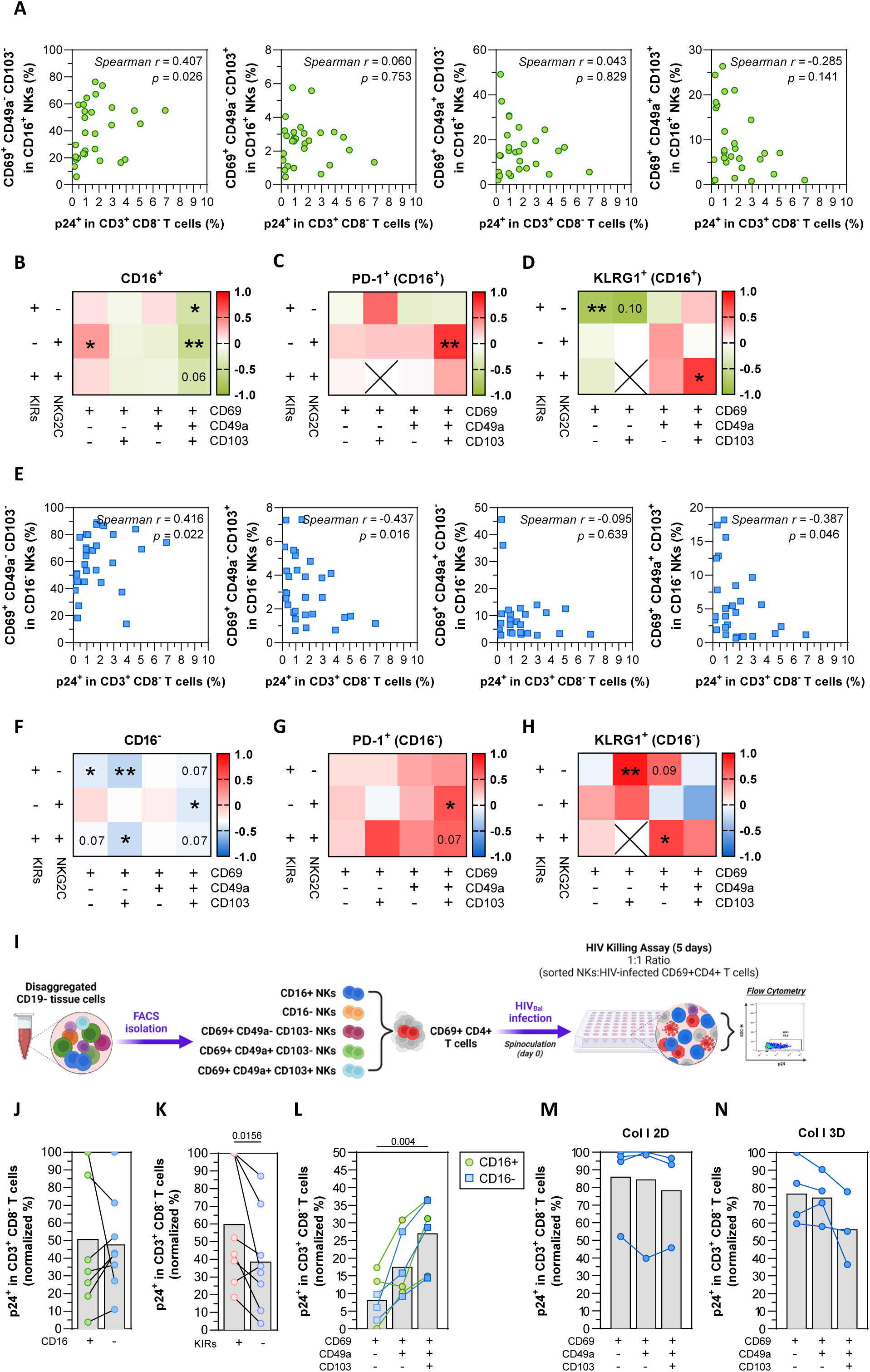
Tonsil-resident NK subpopulations have different capabilities to restrain HIV infection. Tonsillar tissue explants were cultured on hemostatic sponges and *ex vivo*-infected by HIV_BaL_ for 5 days. The expression of resident memory markers and immune checkpoints was measured by flow cytometry at baseline and correlated with the *ex vivo* HIV infection levels (% p24^+^ cells within CD3^+^CD8^−^ T cells). Spearman correlations of the significant results are shown. **(A-H)** Spearman correlations and correlation matrices showing the relationship between HIV p24 levels in explants and the initial frequencies of **(A)** resident CD16^+^, **(B)** resident memory CD16^+^, **(C)** resident memory CD16^+^ PD-1^+^, **(D)** resident memory CD16^+^ KLRG1^+^, **(E)** resident CD16^−^, **(F)** resident memory CD16^−^, **(G)** resident memory CD16^−^PD-1^+^, and **(H)** resident memory CD16^−^KLRG1^+^. **(I)** Experimental setting of functional cocultures to assess the suppression capacity of HIV infection by tonsil-resident NK subpopulations. NK cells were FACS-isolated based on the expression of resident and memory markers and cocultured with autologous CD4^+^CD69^+^ T cells infected by spinoculation with HIV_BaL_. Direct cell killing of HIV^+^CD3^+^CD8^−^ T cells (p24^+^) was quantified by flow cytometry 5 days after infection. **(J-N)** NK cytotoxicity assays of *ex vivo* HIV-infected CD4^+^CD69^+^ T cells after coculturing with isolated NK cells showing differential expression of **(J)** CD16, **(K)** KIRs, and **(L-N)** resident markers. Assays were conducted under three conditions: **(J-L)** suspension culture, **(M)** on a collagen I-coated surface (2D), and **(N)** embedded within a collagen I gel (3D). In **(J)**, **(K)**, **(L)**, **(M)** and **(N)**, different NK subpopulations and CD4^+^CD69^+^ T cells were isolated by FACS. Statistical comparisons were performed using the Wilcoxon matched-pairs signed rank test or Friedman’s test, as appropriate. Significant comparisons *p*<0.05.

Next, to determine the individual cytotoxic potential of different resident NK cell populations to eliminate HIV infected cells, we isolated different tonsillar NK cell populations using FACS based on CD16, CD69, CD49a, CD103 and KIRs expression, and conducted functional killing assays against autologous HIV-infected tonsil-resident CD4^+^ T cells **(Fig 5I)**. Due to the limited availability of NKG2C⁺ NK cells for functional assays, here we could only focus on the KIR receptors. We first assessed the killing capacity of the two main CD56^+^ fraction. No significant differences were observed between both CD16^+^ and CD16^-^ cells (**Fig. 5J**). Next, we assessed the cytotoxic capacity of KIR-expressing cells and found that, in both the CD16⁺ and CD16⁻ fractions, KIR⁺ expression alone does not enhance cytotoxicity. Instead, KIR⁺ cells exhibited a significantly lower ability to kill HIV-infected cells (**Fig. 5K**). This aligns with our previous findings showing that KIR expression is associated with the production of IFN-γ (**Fig. 2J**), a regulatory cytokine rather than one directly involved in cytotoxicity. Finally, we evaluated the impact of individual resident markers on HIV-infected cell killing by establishing three distinct culture conditions: a conventional cell suspension, a collagen I-coated system, and a collagen I-based 3D coculture assay, the latter of which more accurately reflects the in vivo microenvironment and cellular responses to stimuli^67^. Importantly, we observed that NK cell cytotoxicity was strikingly influenced by the culture conditions. In cell suspension, SP NK cells demonstrated significantly superior killing of HIV-infected cells compared to TP NK cells, which exhibited poor control of HIV-infected tonsillar CD4^+^ T cells (92.1% and 70.7% reduction, respectively, *p*=0.004) **(Fig 5L)**. However, on a collagen I-coated 2D surface, the DP and TP populations reached the same killing capacity as the SP population, and no significant differences in killing efficiency were observed among the three sorted resident NK populations during coculture (**Fig. 5M**). In contrast, when NK and HIV-infected cells were embedded in a collagen I gel (3D), TP NK cells demonstrated superior reduction of HIV-infected cells **(Fig 5N)**.

In summary, we found that TP tonsillar NK cells expressing NKG2C are linked to reduced levels of HIV infection, with the expression of immune checkpoint markers significantly correlating with the HIV burden. Additionally, the tissue microstructure emerges as a critical factor in shaping the functionality of resident NK cells. When evaluated outside their native tissue context, cells display a markedly different capacity for HIV control compared to their behavior within the tissue microstructure. These findings highlight the importance of employing advanced models that incorporate the extracellular matrix to accurately evaluate the efficacy of tissue-resident NK cells in cell targeting.

### Impact of ex vivo HIV infection in tonsillar ILCs

We investigated whether HIV infection induces alterations in the phenotype and function of NK cells, focusing on the populations identified as important in responses against both K562 cells and HIV. To this end, we infected tissue blocks and, after 5–6 days, measured HIV replication and characterized tonsillar NK cells using single-cell RNA sequencing (scRNA-seq) analyses and functional assays. After quality control (see *‘Methods’*), 3,562 CD56^+^ cells from uninfected explants and 4,433 CD56^+^ cells from HIV-infected explants were successfully sequenced. Uniform manifold approximation and projection (UMAP)^68^ revealed 11 distinct cell clusters within the ILC lineage across both experimental conditions, with no significant differences in cluster frequency **(Fig. 6A)**. Clusters were manually annotated based on the top 10 highly differentially expressed genes **(Fig. S5)** and defining markers from datasets of fresh, uninfected tonsils **(Fig. S6)**^34,69–73^. Cluster 9 and 11 represented NK cells expressing CD16 (*FCGR3A*).

**Figure 6.**
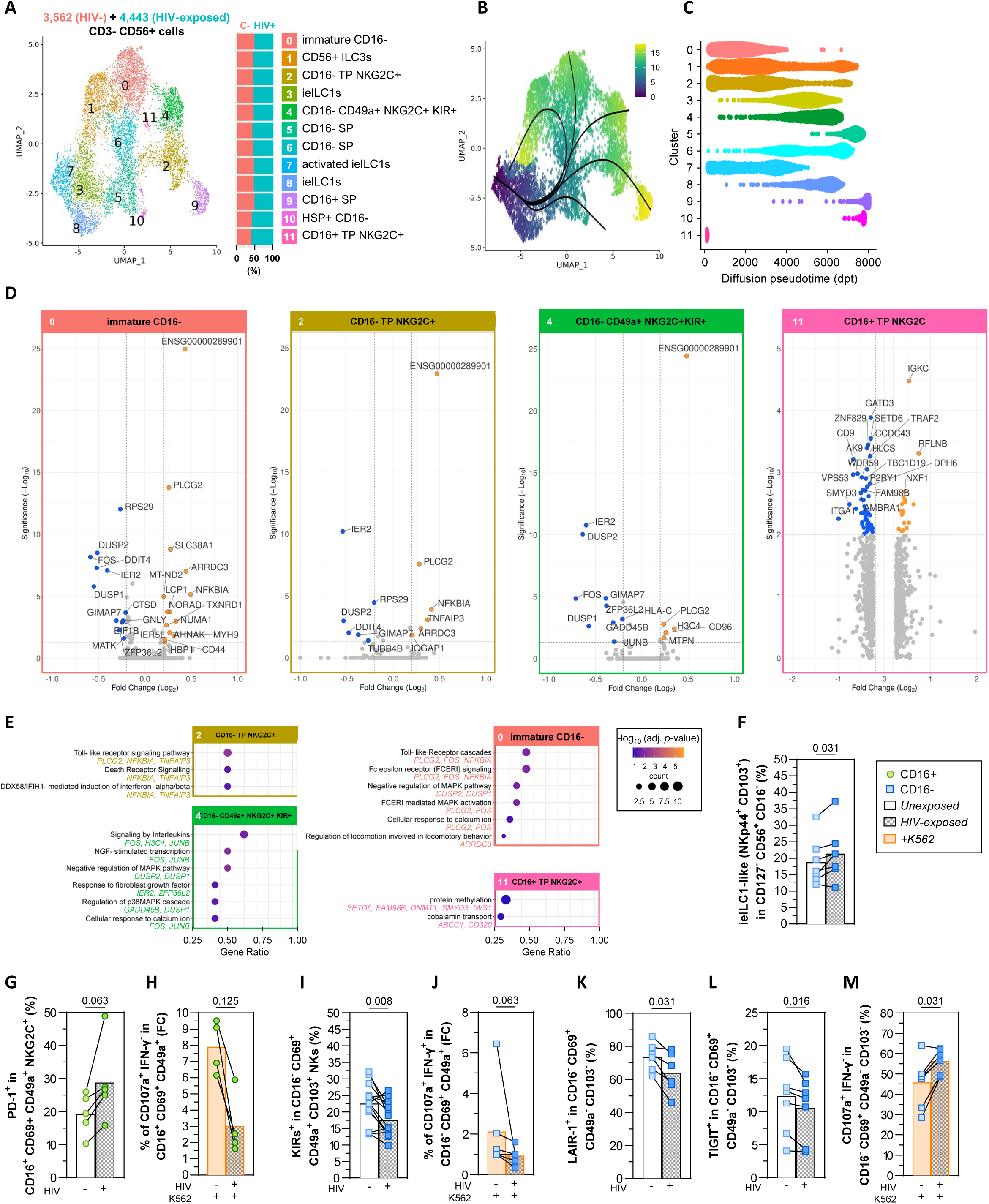
Distinct transcriptional signatures and functional differences in tonsil-resident ILC/NK cells during early HIV exposure. Tonsillar tissue explants were cultured on hemostatic sponges and *ex vivo*-infected by HIV_Bal_ for 5-6 days. The CD14^−^CD19^−^CD3^−^CD56^+^ cell population from HIV-uninfected and HIV-infected explants was isolated by FACS and analyzed using scRNA-seq. **(A)** UMAP visualization of 12 distinct cell clusters identified by unsupervised hierarchical clustering and proportion of cells per cluster and sample (right). **(B)** UMAP representation of diffusion pseudotime trajectories, rooted in activated ieILC1s (cluster 7), indicating maturation states from dark blue (least mature) to yellow (most mature). **(C)** Bee swarm plot illustrating the distribution of cells from the 12 ILC clusters across pseudotime. **(D)** Volcano plots of clusters 0, 2, 4, and 11 showing DEGs between uninfected and infected tissue explant CD56^+^ cells. For cluster 11, note that the volcano plot was generated using unadjusted *p*-values due to its low frequency representation. **(E)** Gene Ontology (GO) and Reactome pathway analyses highlighting selected pathways enriched in clusters 0, 2, 4, and 11 in response to HIV exposure. DEGs represented in each enriched pathway are shown. **(F)** Frequency of ieILC1s in the infected culture compared to the uninfected control. The expression of resident memory markers and immune checkpoints was measured by flow cytometry in the steady state and upon stimulation with K562 cell coculture from uninfected and infected tissue explant cells; **(G)** Expression of PD-1 on CD16^+^CD69^+^CD49a^+^NKG2C^+^ NK cells in the infected culture compared to the uninfected control. **(H)** Degranulation potential (fold-change of CD107a^+^IFN-γ^−^ cells) of CD16^+^CD69^+^CD49a^+^ NK cells in uninfected and infected explants upon K562 stimulation. **(I)** Expression of KIRs on CD16^−^CD69^+^CD49a^+^CD103^+^ NK cells in the infected culture compared to the uninfected control. **(J)** Cytokine production potential (fold-change of CD107a^+^IFN-γ^+^ cells) of CD16^−^CD69^+^CD49a^+^ NK cells in uninfected and infected explants upon K562 stimulation. **(K)** Expression of LAIR-1 on CD16^−^CD69^+^CD49a^−^CD103^−^ NK cells in the infected culture compared to the uninfected control. **(L)** Expression of TIGIT on CD16^−^CD69^+^CD49a^−^CD103^−^ NK cells in the infected culture compared to the uninfected control. **(K)** Degranulation potential (fold-change of CD107a^+^IFN-γ^−^ cells) of CD16^−^CD69^+^CD49a^−^CD103^−^ NK cells in uninfected and infected explants upon K562 stimulation. Statistical comparisons were performed using the Wilcoxon matched-pairs signed rank test. Significant comparisons *p*<0.05. Green circles: CD16^+^; Blue squares: CD16^−^.

We first focused on the deep characterization of cells identified as important in responses against HIV and K562 cells. Cluster 11 was identified as CD16⁺TP NKG2C⁺ cells, and cluster 2 as CD16⁻ TP NKG2C⁺ cells, both of them were highly correlated with lower levels of HIV infection in tissue explants **(Fig. 5B, F)**. Based on the transcriptional data, these NK cells exhibit cytotoxic activity through the expression of molecules like granulysin (*GNLY*), perforin (*PRF1*), and granzymes (*GZMB*) **(Fig. S6)**. They also express molecules involved in the apoptosis of target cells via Fas ligand (*FASLG*) and TRAIL (*TNFS10*). Additionally, they may respond to inflammatory and immunoregulatory cytokines (*IL2RB*, *IL4R*, *IL10RA*) and express receptors involved in migration to infection sites via *CCL3-5*, *CCR7*, and *SELL*. These cells also express molecules that modulate immune responses through *TGFBR3* and *PTGER2-4* **(Fig. S6)**. Overall, these characteristics suggest that these populations have a prominent role in cytotoxicity, inflammation, and immune modulation. Interestingly, these clusters also express NKG2A (*KLRC1*), and a similar cell population in the blood has been associated with HIV control in elite controllers—individuals living with HIV who do not require antiretroviral therapy to control the infection^74^.

Next, we identified the CD16⁻CD49a⁺KIR^+^ population as cluster 4, which exhibited the highest levels of CD107a and IFN-γ production following K562 stimulation **(Fig. 2J)**. This population closely resembled the previous cluster 2, but with a distinctive expression profile of inhibitory KIRs (*KIR2DL1*, *KIR2DL3*, *KIR3DL1*, *KIR3DL2*, *KIR3DL3*) and the activating *KIR2DS4*, in addition to NKG2C (*KLRC2*) **(Fig. S6)**. This suggests that while cluster 4 is highly functional in terms of cytotoxicity and cytokine production, its activity may be modulated by the balance of KIR receptors.

Other ILCs include: cluster 1 identified as CD56^+^NKp44^+^ ILC3s (enrichment of *KIT*, *IL1R1*, *IL23R*, *RORC*, *CD300LF* and *NCR2)*^70^. Clusters 3, 7, and 8 corresponded to ieILC1s, distinguished by their expression of NKp44 (*NCR2*) and CD103 (*ITGAE*)^52^. Cluster 7 was classified as activated ieILC1s due to its highest expression of *CD69* and *IL27RA*. Cluster 0 represented immature CD16⁻CD69⁺ NK cells and was marked by the expression of NKG2A (*KLRC1*) and homing markers for secondary lymphoid organs (*CCR7* and *SELL*)^75^. Additional CD16⁻CD69⁺ NK cells were represented by clusters 5 and 6, with only minor differences between the two. Finally, cluster 10 comprised NK cells with high expression of heat-shock proteins (HSP) and stress-response genes like *DNAJB1*^72^ **(Fig. S6)**.

Next, using pseudotime inference analysis^76^, we traced the differentiation trajectories of tonsil-resident ILCs within tissue explants. Activated ieILC1s were identified as the common ancestor, transitioning through CD16⁻CD69⁺ NK cells and exhibiting plasticity by diversifying into both distinct terminal states and immature subsets **(Fig. 6B)**. Diffusion map pseudotime highlighted clusters 5 and 6 (CD16⁻CD69⁺), 9 (cytotoxic CD16⁺), and 10 (HSP+ CD16⁻) as terminally enriched **(Fig. 6C)**, indicating maturation within the tonsils and a likely readiness for circulation^37^. Notably, clusters 5, 6, and 9 showed the lowest mRNA expression of *RGS1* **(Fig. S6)**, a key marker of tissue-infiltrating lymphocytes^73^. Interestingly, cluster 11 (CD6⁺TP NKG2C⁺) exhibits a short pseudotime, which might indicate either rapid differentiation without transitioning through multiple intermediate states or an alternative differentiation pathway.

After analyzing ILC populations in tonsils, we aimed to dissect the impact of HIV infection on these cells. To achieve this, we performed differential gene expression (DEG) analysis, which highlighted key transcriptional changes induced by HIV exposure. A comprehensive list of DEGs per cluster is provided in **Supplementary Table S2,** and main changes within selected clusters were represented in volcano plots **(Fig. 6D and S7A)**. Additionally, Reactome and Gene Ontology term enrichment analyses were performed to infer altered pathways in NK cell clusters after HIV exposure **(Fig 6E and S7B)**. Interestingly, the lncRNA *ENSG00000289901* was upregulated across clusters 1-9 following HIV exposure **(Fig. S8A)**. Using CATrapid omics v2.1, we identified top predicted protein interactions **(Fig. S8B)**, including those directly involved in the regulation of HIV replication (ZFC3H1^77^ and HTATSF1^78^). However, the function of this lncRNA is currently unknown.

HIV infection induced significant gene expression changes in NK cells from the functional clusters 2, 4 and 11. In cluster 11 key downregulated genes included *CD9*^79^, which is involved in cell adhesion and the formation of the immune synapse, likely impairing the interaction with HIV-infected targets, *TRAF2*^80^, reducing TNF receptor signaling and weakening inflammatory responses, and *ITGA1*, limiting NK cell migration and tissue infiltration. Additionally, *SETD6*^81^ and *SMYD3*^82^ affect epigenetic regulation, likely suppressing NK cell activation, while *GATD3*^83^ and *AK9*^84^ disrupt metabolic pathways essential for energy production and proliferation (**Fig 6D, E**). For cluster 2 HIV infection introduced changes compatible with impaired NK antiviral functions. Downregulated genes include *IER2* (modulating adhesion to collagen type I^85^), *RPS29* (limiting protein synthesis for cytotoxic function^86^), *DUSP2* (disrupting immune signaling^87^), and *DDIT4* (weakening metabolic fitness^88^). *GIMAP7*^89^ and *TUBB4B*^90^, involved in cell survival and cytoskeletal dynamics, are also reduced, likely impairing NK cell persistence and target engagement. Conversely, upregulated genes like *PLCG2* (enhancing abnormal degranulation^91^), *NFKBIA* and *TNFAIP3*^92^ (suppressing NF-κB-mediated immune responses), *ARRDC3* (affecting integrin trafficking^93^), and *IQGAP1* (disrupting cytoskeletal organization^94^) might promote NK cell dysfunction (**Fig 6D, E**). HIV also disrupted genes regulating IFN-γ production, particularly in cluster 4 (defined previously as the most functional subset responding to K562 cells, characterized as CD16⁻CD69^+^CD49a^+^KIRs^+^) **(Fig. 2I)**. They downregulated *GADD45B, FOS, JUNB, DUSP1, DUSP2,* and *ZFP36L2* likely impairing key immune functions. GADD45B is critical for IFN-γ production in tumor-specific CD8^+^ T cells^95^ and for full activation of CD56^bright^ NK cells^96^, and regulates stress responses and apoptosis^97^, while FOS and JUNB drive immune activation and cytokine production^98,99^. DUSP1 and DUSP2 modulate MAPK signaling^87^, and ZFP36L2 controls immune gene expression, with their loss leading to dysregulated responses^100^. Simultaneously, NK cells upregulated CD96 and AHNAK, where CD96 functions as an inhibitory receptor dampening cytotoxicity^101,102^, and AHNAK influences migration and adhesion^103^ (**Fig 6D, E**). These changes suggest that HIV induces NK cell dysfunction by impairing activation, disrupting signaling, and increasing inhibitory regulation, likely weakening antiviral responses in the most activated subsets.

Other important changes following HIV exposure in the remaining clusters include the immature CD16⁻ NK cells (cluster 0). Although they share some altered genes with cluster 2, we also observed the exclusive upregulation of genes suggesting a shift in NK cell functionality towards a more energetically prepared state, potentially indicating functional reprogramming. The upregulation of genes like *SLC38A1*^104^ suggests enhanced signaling and metabolic activity. Additionally, upregulation of mitochondrial and metabolic genes (*MT-ND2*, *TXNRD1*^105^) points to increased energy production, while changes in adhesion molecules (*CD44*^106^), increased motility (*LCP1*)^107^, cell proliferation (*NUMAI*)^108^, and cytoskeletal components (*MYH9*^109^) might suggest readiness for migration and tissue-specific responses (**Fig 6D, E**). Overall, this gene expression profile might indicate that NK cells are likely primed for a more efficient immune response. In contrast, we observed that HIV exposure introduced changes in the transcriptome of cluster 1 (CD56^+^ ILC3s) and 7 (activated ieILC1s) compatible with impaired IL-15 signaling, as evidenced by the downregulation of *ID2*, which suppresses SOCS3 expression to maintain IL-15 signaling^110^. Furthermore, cluster 7 exhibited pathways altered by HIV exposure included Rho GTPase effectors, cellular senescence, I-kappaB/NF-kB signaling, tissue remodeling, and epigenetic changes (histone H3-K4 methylation and histone H3-K27 trimethylation) (**Fig. S7A, B**).

In conclusion, HIV exposure induces rapid transcriptomic changes in tonsillar NK cells, including increased migration and proliferation potential in immature NK cells (cluster 0), tightly regulated IFN-γ production in resident memory NK cells (cluster 4), while impairing the killing capacity of more cytotoxic and differentiated NK cells (clusters 2, and 11). These findings suggest that resident NK cell subsets undergo rapid functional adaptation to early HIV infection in tonsillar tissue.

To complement previous results, we evaluated the impact of HIV infection on selected NK cell populations performing functional assay with K562 cells. Our results did not show a significant global redistribution of the tonsil-resident CD56^+^ NK cell compartment in response to HIV infection **(Fig. S9A-D)**, which is consistent with the single-cell transcriptomic findings. However, ieILC1s were slightly expanded upon HIV infection **(Fig. 6F)** and certain subpopulations exhibited significant variation in the expression of memory markers, ICs, and functional markers after K562 coculture. In the CD16+ fraction, we observed higher expression of PD-1 on the cytotoxic CD16^+^CD69^+^CD49a^+^ NK cells after HIV infection **(Fig. 6J)**, along with reduced degranulation upon K562 stimulation **(Fig. 6H)**. In the CD16⁻ population, KIR expression was reduced in the CD49a^+^CD103^+^ fraction **(Fig. 6I)**, a marker that we identified as important for the production of CD107a and IFN-γ upon cell encounters. Accordingly, the production of both functional markers was diminished **(Fig. 6J)**. Furthermore, when evaluating the maximum stimulatory potential of more immature CD16^−^SP, we found that HIV infection led to downregulation of inhibitory receptors LAIR-1 **(Fig. 6K)** and TIGIT **(Fig. 6L)**. This suggests a functional shift that could enhance their activation, leading to an increase in CD107a production upon target cell stimulation **(Fig. 6M)**. These findings align with the previous transcriptome results, reinforcing the idea that HIV infection impairs NK cell cytotoxicity in more differentiated cells while enhancing function in less mature populations.

## DISCUSSION

The mechanisms underlying the rapid mounting of local NK cell responses to initial HIV infection in secondary lymphoid tissues are still poorly understood. Despite the innate nature of NK cells^111^, circulating NK cells have shown immunological memory to certain viruses^40^, including HIV^42^. In tissues, resident NK subpopulations respond firmly to secondary infections^25,29,44,112^ and possess recall responses in vaccinated individuals^29,44^. Here, we investigated the functional diversity of tonsil-resident NK cells and their anti-HIV activity. We observed that specific NK subsets exhibited cytotoxic and antiviral effects. When NK cells were cultured with K562 targets, the CD16⁺ TP cells showed enhanced degranulation, whereas the CD16⁻ DP KIR⁺ cells produced the most IFN-γ. Notably, the TP NKG2C subset (both CD16^−^ and CD16^+^) correlated with lower HIV infection levels and was the most effective at killing HIV-infected cells in extracellular matrix-based functional assays. However, HIV profoundly impaired the functionality of these antiviral NK cells, while inducing changes in more immature populations, consistent with immune reprogramming.

We observed that despite being less frequent in the CD16^−^ fraction, CD49a^+^KIRs^+^ NK cells (DP) show heightened IFN-γ production when stimulated with K562 cells, suggesting functional differences linked to CD16 and CD49a expression. Functional assays reveal that CD49a enhances NK cell activity, likely by improving their ability to interact with target cells through the tissue extracellular matrix^113,114^. Of particular interest is the observation that IFN-γ in tissues inhibits collagen synthesis^115,116^, thereby facilitating access for killer lymphocytes to localized infections. Additionally, some direct effects of IFN-γ-producing NK cells towards the extracellular matrix have been identified^117^. Our results are consistent with previous studies; in humans, there is compelling evidence that CD16^−^CD49a^+^ NK cells display higher production of IFN-γ^32^. Interestingly, despite CD49a^+^ NK cells being proinflammatory across human tissues, certain conditions can exacerbate their cytotoxic status. In the human lung, they were found to produce perforin and granzyme B at levels comparable to the CD49a^−^fraction when stimulated with IL-15^31^. Moreover, CD16^−^CD49a^+^ NK cells from the center of lung tumors degranulated stronger than CD16^−^CD49a^−^ NK cells^63^. Indeed, recall responses have been attributed to these resident NK cells for Varicella-Zoster Virus^29^ and Hepatitis A and B viruses^44^. Of note, a strong parallelism with tissue-resident memory CD8^+^ T cells (trm CD8^+^ T cells) exists. After influenza infection, lung trm CD49a^+^CD8^+^ T cells slowly upregulate mRNA levels of perforin, granzymes and death-receptor ligands^118^, reinforcing the idea that CD49a^+^ lymphocytes act as initial producers of antiviral IFN-γ against a first-ever-seen pathogen, serving as a warning system, and enhance their cytotoxic response when the infection persists longer^25,44,119,120^.

After mapping the functionality of tonsil-resident NK cells against the conventional targets K562 cells, we used the tonsillar explant model to define the effect of HIV infection on the natural NK response. Importantly, CD103 expression on NK cells was strongly correlated with HIV control in explants, with these CD16⁻CD103⁺ cells being part of the intraepithelial innate lymphoid cell (ieILC) compartment. Interestingly, the expression of CD103 is guided by TGF-β ^121^, an immunosupressive cytokine with known effects on NK cells^62^. However, CD103^+^ NKs may be crucial to control epithelial infections since these NKs localize intraepithelially^49^. During the resolution of the ongoing infection, immunosuppressive TGF-β increases CD103 in CD49a^+^ lymphocytes^122,123^ to cross basement membrane by granzyme B liberation^124^ and transmigrate to establish intraepithelial residence, most likely to combat new infections entering from epithelial barrier. Thus, while TGF-β induces CD103 as part of an immunoregulatory mechanism, it may also contribute to antiviral defense by guiding NK cells into epithelial tissues. This raises the possibility that harnessing CD103⁺ NK cells could enhance mucosal immunity and inform strategies to limit HIV persistence and transmission. Interestingly, we observed that these CD103⁺ NK cells exhibited superior cytotoxicity against HIV-infected cells in a 3D collagen gel coculture system, a phenomenon not observed in suspension cultures. Collagen I (Col I), a major component of the tissue interstitial matrix, plays a crucial role in supporting the migration of cytotoxic lymphocytes^114^, which likely explains the enhanced cytotoxic advantage of CD103⁺ NK cells in this context. However, a negative effect is the promotion of tissue HIV infection by cell-to-cell transmission^125,126^. These findings emphasize the critical importance of studying functional NK cell responses within their appropriate microenvironment. For example, IL-15, which can be transpresented by dendritic cells in the tonsils^37,127^, upregulates cytotoxic mediators in CD16^−^CD69^+^CD49a^+^ NK cells^31,44,128^ and counteracts the immunosuppressive effects of TGF-β^62,129^, thereby promoting the optimal response of ieILC1s^52^. Of note, chronic HIV infection leads to TGF-β-mediated collagen deposition that exacerbates tissue compartmentalization^13,14,130^.

While it is well established that HIV impairs NK cell function^131^, most studies to date have focused on peripheral blood, leaving the impact on tissue-resident NK cells largely unexplored. Here, we provide evidence that HIV-driven NK cell dysfunction extends to tissue compartments, identifying the specific subsets affected. Our findings demonstrate that HIV infection profoundly impacts the functionality of tissue-resident NK cells, driving significant phenotypic and functional changes. Specifically, we observed increased expression of PD-1 and KIRs in more differentiated, antiviral NK cell subsets, aligning with previous reports in blood showing elevated immune checkpoints and KIR expression in NK cells from viremic individuals^19,131,132^. Moreover, we analyzed the transcriptomic profile of these cells and confirmed the downregulation of genes associated with cell activation and cytotoxic capacity, such as *CD9, FOS, JUNB, GADD45B, DUSP1-2 and NKKB1A*. In the context of viral infections, CD9 may influence immune responses by regulating leukocyte adhesion and migration, thereby affecting the recruitment of immune cells to sites of infection^79^. However, the specific impact of CD9 upregulation on NK cell cytotoxicity, especially in the context of HIV infection, requires further investigation. Moreover, *GADD45B* is critical for IFN-γ production in tumor-specific CD8^+^ T cells^95^ and for full activation of CD56^bright^ NK cells^96^. Key components of the AP-1 complex (*FOS* and *JUNB*) were also downregulated. Moreover, *DUSP1* and *DUSP2*, which inactivate MAP kinases that act upstream in AP-1 activation^87^, were downregulated, suggesting that AP-1 signaling, essential for IFN-γ production^133^, is impaired at multiple points in the signaling cascade. Furthermore, the loss of *DUSP1* transcription increases IFN-γ and CD107a expression^87^, and *ZFP36L2* downregulation, an IFN-γ translation inhibitor during prolonged T cell activation^100^, further supports an adaptive mechanism to preserve IFN-γ production despite persistent HIV-driven activation. These cells also increased *CD96* expression, encoding for an immune checkpoint linked to intratumoral NK cell exhaustion^101,102^, and upregulated *AHNAK* suggest TGF-β may drive NK dysfunction. Notably, TGF-β blockade restores CD96^+^ NK cell functionality^101^. Overall, these findings highlight that HIV induced dysregulation of critical signaling pathways in activated and cytotoxic clusters, alongside alterations in immune checkpoints and adaptive mechanisms, collectively contribute to impaired IFN-γ production and NK cell dysfunction, while also revealing potential targets such as TGF-β for restoring immune functionality.

Conversely, we observed that HIV infection induced a state of hyperresponsiveness in the more immature CD16^−^CD69^+^ NK cell population within tonsil explants. Notably, HIV not only activated a subset of CD16^−^ tissue-resident NK cells but also heightened their reactivity to subsequent target cell stimulation. This enhanced responsiveness coincided with a marked downregulation of inhibitory receptors, including LAIR-1 and TIGIT. These changes were particularly pronounced in the CD69^+^ SP NK cell subset, which exhibited a striking shift toward a more activated and responsive state. Intriguingly, CD69 expression alone has been linked to the acquisition of a resident memory phenotype, as demonstrated in a murine model of skin infection^25^, suggesting a potential parallel in the functional reprogramming of NK cells during HIV infection. Moreover, receptors such as TIGIT are found at higher levels on NK cells from HIV-infected individuals compared to healthy controls. This upregulation is associated with decreased production of IFN-γ and impaired cytotoxic responses, indicating a state of NK cell exhaustion^134^. While specific studies on LAIR-1 expression in HIV-infected individuals are limited, LAIR-1 is recognized as a relevant receptor in the context of NK cell regulation and immune checkpoints^135^. Therefore, the observation of enhanced NK cell responsiveness coinciding with a marked downregulation of inhibitory receptors, such as LAIR-1 and TIGIT, contrasts with the typical upregulation of these receptors during HIV infection. This suggests a complex interplay between NK cell activation and inhibition, which may vary depending on the stage of infection, treatment status, tissue compartment or other immunological factors. Furthermore, and consistent with this functional reprogramming, we observed transcriptional changes indicative of enhanced activity, including the upregulation of genes such as *PLCG2*, essential for NK cell-mediated antiviral responses^91^, *SLC38A1,* vital for energy demand of activated lymphocytes^136^, *MT-ND2,* involved in NK cell metabolism^137^, and *NUMA1,* involved in cell division and proliferation^138^. Importantly, we also identified the upregulation of genes associated with increased NK cytotoxicity and migration, such as *MYH9*^109^ and *CD44*^106^, further supporting the transition to a more tissue-adapted and functionally dynamic phenotype. Importantly, CD44 interacts with Galectin-9, a β-galactoside-binding lectin. This interaction has been shown to promote NK cell activity, enhancing their effector functions. Notably, increased Galectin-9 expression has been observed in certain viral infections, including HIV^139^, suggesting a role for the Galectin-9/CD44 axis in modulating NK cell responses during infection^140^.

In conclusion, our study provides novel insights into the phenotypical and functional characteristics of human NK cells within secondary lymphoid tissues, both under steady-state conditions and during the early stages of HIV infection. Our findings highlight the significance of considering specific spatial and immunological factors in the design of future therapeutic interventions targeting HIV reservoirs within tissues, with a focus on harnessing the potential of tissue-resident memory NK cells.

### LIMITATIONS OF THE STUDY

This study using the tonsillar tissue model predominantly focuses on changes occurring within the local NK cell populations, without fully considering potential contributions from NK cells recruited from circulation. This is noteworthy given that a significant portion of NK cells in tonsils consists of immature subpopulations undergoing differentiation before they enter circulation^37,53,141^. Nonetheless, the tonsillar explant model presents a unique opportunity to focus exclusively on NK cells that reside within the tonsils, independent of those in circulation. Moreover, tissue remodeling by matrix metalloproteinases (MMPs) results in the shedding of CD16^142^, introducing the likelihood that functional (CD107a^high^) CD16^−^ SP might have initially been CD16^+^ SP NK cells. Prior activation could induce degranulation and CD16 shedding, a process known to be advantageous for promoting serial killing^143^. Finally, our focus on CD49a and CD103 expression may oversimplify the diversity within resident NK cells. Moreover, coexpression of both CD49a and CD49b identifies currently active NK cells, while those expressing only CD49a are considered long-term residents^144,145^. Addressing these limitations will strengthen the robustness and applicability of our findings in the broader context of immune responses to tissue infections.

## Materials and Methods

### Patients samples and ethics statement

In this study, we used primary human tissue cells recovered from tonsillar resections obtained from routine surgery for treating chronic snoring or sleep apnea in children under 18 years old. Tissue samples were obtained from the Otorhinolaryngology Unit of the Hospital Universitari Vall d’Hebron in Barcelona, Spain. Study protocols were approved by the corresponding Ethical Committees (Institutional Review Board numbers PR(AG)270/2015 and PR(AG)582/2020). All subjects recruited for this study were children under 18 years old who provided parental written informed consent. Information on gender, age, and cause of surgery is summarized in **Supplementary Table S1**. As we used samples from donors of different gender, we might conclude that findings on the functionality of trNK cells to eliminate the HIV reservoir apply for both, men and women.

### Culture human tissue explants

Tonsillar specimens were processed in fresh upon collection for phenotypical studies or let overnight in dissection medium when intended for culturing or functional assays to ensure the inactivation of contaminating microorganisms. Dissection medium included RPMI-1640 medium (Gibco) supplemented with 500 U/ml penicillin (Fisher Scientific), 500 µg/ml streptomycin (Fisher Scientific), 1 µg/ml gentamicin (Gibco), and 5 µg/ml amphotericin B (Gibco). Tonsillar tissue explants (5×5 mm blocks) were cultured on the air-liquid interface on absorbable gelatin sponges (Goodwill) as previously described^66^. The hemostatic sponges were soaked in RPMI-1640 medium supplemented with 20% Fetal Bovine Serum (FBS) (Gibco), 10 mM HEPES (Biowest), 100 U/ml penicillin, 100 µg/ml streptomycin, and 66 µg/ml amikacin (Vall d’Hebron Hospital pharmacy) (R20 medium). The first day of culture, R20 medium also included 310 µg/ml of timentin (Caisson Labs). For the culture of digested and sorted tonsillar tissue cells, we used R20 medium supplemented with 100 U/ml interleukin-2 (IL-2) obtained from the Vall d’Hebron Hospital pharmacy. In all the experiments, medium was changed every 2-3 days and cultures were maintained at 37°C in a 5% CO_2_ incubator. In order to infect tonsillar tissue explants, we deposited 457 TCID_50_µl of HIV_BaL_ on each tissue block cultured on sponges. For infection of sorted tonsillar CD4^+^CD69^+^ T cells and Dynabeads-isolated CD4^+^ T cells, we employed a spinoculation method^146^ in which 300,000 target cells were centrifuged in a 96-well conical bottom plate with a 50% tissue culture infectious dose (TCID_50_) of 625 units of HIV_BaL_ at 1,200 x *g* for 2 h at 37 °C. Then, cells were washed twice with Dulbecco’s PBS 1X (Capricorn Scientific) to remove unbound virus. Uninfected cells were spinoculated with R10 alone and served as infection controls.

To obtain the suspension of tonsillar cells, tonsillar resections were minced into small pieces (5×5 mm) avoiding cauterized and excessively bloody, fatty and pus-containing parts. 10 blocks were added per 1.5 ml Eppendorf containing 400 µl of RPMI-1640 medium supplemented with 5% FBS (R5 medium) and were mechanically digested using pestles (Labolan) coupled to a handheld homogenizer (Labolan). Tonsillar cell suspension was subsequently filtered through 70 and 40 µm pore size cell strainers (LabClinics), and 30 µm pore size CellTrics® filters (Sysmex). Filtered cells were washed twice with PBS 1X and the remaining red blood cells were lysed using ACK Lysis Buffer (Fisher Scientific). Cell number and viability were assessed with LUNA Automated Cell Counter (Logos Biosystems).

### Tonsillar resident memory NK cell phenotyping

Fresh tonsillar tissue was digested, and the resulting cell suspension was stained with LIVE/DEAD AQUA viability (ThermoFisher) for 20 minutes at room temperature (RT). After washing once with PBS 1X, cells were stained with anti-CD56-FITC (B159; Becton Dickinson), anti-CD49a-PE-Cy7 (TS2/7; BioLegend), anti-pan-KIRs (KIRs2DL1/S1 and KIRs2DL2/3/S2)-PE-Cy5.5 (EB6B & GL183; Beckman Coulter), anti-CD69-PE-CF594 (FN50; Becton Dickinson), anti-NKG2C-PE (134591; R&D Systems), anti-CD3-APC (SK7; BioLegend), anti-CD16-BV786 (3G8; Becton Dickinson), anti-CD103-BV650 (Ber-ACT8; Becton Dickinson), anti-CD45-BV605 (HI30; Becton Dickinson), anti-CD14-V500 (MφP9; Becton Dickinson), and anti-CD19-V500 (HIB19; Becton Dickinson) antibodies mixed with Staining Buffer (PBS 1X 3% FBS) containing 50% Brilliant Stain Buffer (Becton Dickinson) for 20 minutes at RT. Cells were then washed with Staining Buffer (SB) and were fixed with 2% PFA. Samples were acquired on a BD LSR Fortessa flow cytometer and data were analyzed using OMIQ software.

### Tonsillar innate lymphoid cell phenotyping

Tonsillar cell suspension was assessed by flow cytometry to phenotype innate lymphoid cells (ILCs) and NK cell populations. Cells were stained with LIVE/DEAD AQUA viability (ThermoFisher) for 20 minutes at RT. After washing once with PBS 1X, cells were stained with anti-c-kit-BB700 (YB5.B8; Becton Dickinson), anti-CD56-FITC (B159; Becton Dickinson), anti-CD49a-PE-Cy7 (TS2/7; BioLegend), anti-pan-KIRs (KIRs2DL1/S1 and KIRs2DL2/3/S2)-PE-Cy5.5 (EB6B & GL183; Beckman Coulter), anti-CD69-PE-CF594 (FN50; Becton Dickinson), anti-NKG2C-PE (134591; R&D Systems), anti-CRTH2-APC-Vio770 (REA598; Miltenyi), anti-CD45-AF700 (HI30; BioLegend), anti-CD3-APC (SK7; BioLegend), anti-CD16-BV786 (3G8; Becton Dickinson), anti-CD103-BV650 (Ber-ACT8; Becton Dickinson), anti-CD127-BV605 (A019D5; BioLegend), anti-CD14-V500 (MφP9; Becton Dickinson), anti-CD19-V500 (HIB19; Becton Dickinson), and anti-NKp44-BV421 (p44-8; Becton Dickinson) antibodies mixed with SB containing 50% Brilliant Stain Buffer (Becton Dickinson) for 20 minutes at RT. Cells were then washed with SB and were fixed with 2% PFA. Samples were acquired on a BD LSR Fortessa flow cytometer and data were analyzed using OMIQ software.

### Immune checkpoint phenotyping of tonsil-resident memory NK cells

Fresh tonsillar tissue was digested, and the resulting cell suspension was assessed by flow cytometry to measure the expression of immune checkpoints on tonsil-resident memory NK cells. Tonsillar cell suspension was stained with LIVE/DEAD AQUA viability (ThermoFisher) for 20 minutes at RT. After washing once with PBS 1X, cells were stained with anti-CXCR5-BB700 (RF8B2; Becton Dickinson) or anti-CD137-PerCP-Cy5.5 (4B4-1; BioLegend), anti-CD56-FITC (B159; Becton Dickinson), anti-CD49a-PE-Cy7 (TS2/7; BioLegend), anti-pan-KIRs (KIRs2DL1/S1 and KIRs2DL2/3/S2)-PE-Cy5.5 (EB6B & GL183; Beckman Coulter), anti-CD69-PE-CF594 (FN50; Becton Dickinson), anti-NKG2C-PE (134591; R&D Systems), anti-PD-1-APC-Fire750 (EH12.2H7; BioLegend) or CD25-APC-Cy7 (M-A251; Becton Dickinson), anti-CD3-AF700 (SK7; BioLegend), anti-TIGIT-AF647 (MBSA43; ThermoFisher) or anti-NKG2A-APC (Z199; Beckman Coulter), anti-CD16-BV786 (3G8; Becton Dickinson), anti-CD103-BV650 (Ber-ACT8; Becton Dickinson), anti-CD45-BV605 (HI30; Becton Dickinson), anti-CD14-V500 (MφP9; Becton Dickinson), anti-CD19-V500 (HIB19; Becton Dickinson), and anti-LAIR-1-BV421 (DX26; Becton Dickinson) or anti-KLRG1-BV421 (14C2A07; BioLegend) antibodies mixed with SB containing 50% Brilliant Stain Buffer (Becton Dickinson) for 20 minutes at RT. Cells were then washed with SB and were fixed with 2% PFA. Samples were acquired on a BD LSR Fortessa flow cytometer and data were analyzed using OMIQ software.

### NK natural cytotoxicity assay

We used the K562 cell line (Merck), a highly sensitive standard *in vitro* target model to study the natural cytotoxicity function of natural killer cells. These cells were cultured in R10 medium and maintained at 37°C in a 5% CO_2_ incubator. We checked the absence of contaminants in our cell cultures. NK natural cytotoxicity potential was measured by the detection of CD107a and IFN-γ by flow cytometry after coculture with the K562 cells. The tonsillar cell suspension containing resident NK cells was depleted from CD19^+^ cells using a commercial kit (EasySep™ Human CD19 Positive Selection Kit II; StemCell) and cocultured with K562 cells at a 10:1 effector to target ratio (CD19^−^ fraction (containing NK cells): K562) in a 96-well round (U) bottom plate for 4 h at 37 °C and 5% CO_2_. Previous the 4 h coculture, cells were seeded by contact centrifugation at 400 x *g* for 3 min. Anti-CD107a-BV421 antibody (H4A3; Becton Dickinson), BD Golgiplug Protein Transport Inhibitor (Becton Dickinson), and BD GolgiStop Protein Transport Inhibitor containing monensin (Becton Dickinson) were also added to each well at the recommended concentrations. Cells were then washed and stained for cell viability and surface markers as described in the ‘*Tonsillar resident memory NK cell phenotyping*’ Methods’ section. Cells were then fixed and permeabilized with Fixation/Permeabilization Solution (Becton Dickinson) for 20 minutes at 4°C, washed with BD Perm/Wash buffer and stained with anti-IFN-γ-AF700 (B27; Life Technologies) for 30 minutes at RT. After washing, cells were fixed with PFA (2%) and acquired on a BD LSR Fortessa flow cytometer (Becton Dickinson). Data were analyzed using OMIQ software.

### Extracellular matrix effects on the natural cytotoxicity of tonsil-resident NK cells

Effects of extracellular matrix components on resident NK activation were measured by the detection of CD107a and IFN-γ by flow cytometry after coculture with the K562 cells. The tonsillar cell suspension containing resident NK cells was depleted from CD19^+^ cells using a commercial kit (EasySep™ Human CD19 Positive Selection Kit II; StemCell) and NK cells were purified using the CD56^+^ Microbeads (Miltenyi Biotec). Cocultures with K562 cells were established at 2:1 effector to target (NK: K562) ratio in different supports: R10 suspension, 10 µg/cm^2^ collagen (Col) I and IV coating (2D), and 1.5 mg/ml Col I and 1 mg/mL Col IV 3D gels. For the collagen preparations, 96-well flat-bottom plates and 6-well plates were treated with enough collagen solution volume to cover the wells and were left 30 min at 37 °C and 5% CO_2_ for solidification and successful coating. Exceeding volumes were removed and coated wells were washed with warm PBS 1X. For the generation of collagen 3D gels, acidic collagen stock solutions were neutralized with 2M NaOH (Sigma Aldrich) and buffered with HEPES 1M. 3 million NKs/ml and K562 cells were immediately embedded in the collagen gels maintained on ice to prevent rapid solidification before plating or seeded on collagen-coated plates. To investigate the impact of collagen coating on CD49a^+^ NK cell production of IFN-γ, we conducted experiments using a cell density of 70,000 NK cells/cm^2^. This setup ensured adequate spacing between effector and target cells, promoting migratory interactions. Cells were incubated for 16 h at 37 °C and 5% CO_2_. Anti-CD107a-BV421 antibody (H4A3; Becton Dickinson), BD Golgiplug Protein Transport Inhibitor (Becton Dickinson), and BD GolgiStop Protein Transport Inhibitor containing monensin (Becton Dickinson) were also added to each well at the recommended concentrations. After the overnight incubation, cells were treated with collagenase IV (Roche) at 1.4 mg/ml for 30 min at 37 C° and 5% CO_2_ while mixing well content every 10 min to disaggregate the collagen matrices and recover the embedded cells. Cells were then washed and stained for cell viability, surface, and intracellular markers as described in the ‘*NK natural cytotoxicity assay*’ Methods’ section. Stained cells were acquired on a BD LSR Fortessa flow cytometer. Data were analyzed using OMIQ software.

### Generation of viral stock

The plasmid needed for the generation of viral stock were obtained through the NIH AIDS Reagent Program. Viral stocks were generated by transfection of 293T cells (ATCC) with the plasmids encoding the HIV_BaL_, and the resulting viral particles were titrated in TZM-bl cells using an enzyme luminescence assay (Britelite plus Reporter Gene Assay System; Revvity) as described previously^147^.

### FACS isolation of tonsil-resident memory NK cells and killing of autologous HIV-infected cells

Tonsillar tissue was digested, and CD19^−^ cells were purified from the resulting cell suspension using the EasySep™ Human CD19 Positive Selection Kit II. Approximately 100 million CD19^−^ tonsillar cells were stained for Fluorescence-Activated Cell Sorting (FACS) with LIVE/DEAD AQUA viability (ThermoFisher) for 20 minutes at RT. After washing once with PBS 1X, cells were stained with anti-CD56-FITC (B159; Becton Dickinson), anti-CD49a-PE-Cy7 (TS2/7; BioLegend), anti-pan-KIRs (KIRs2DL1/S1 and KIRs2DL2/3/S2)-PE-Cy5.5 (EB6B & GL183; Beckman Coulter), anti-CD69-PE-CF594 (FN50; Becton Dickinson), anti-NKG2C-PE (134591; R&D Systems), anti-CD19-APC-Fire750 (HIB19; BioLegend), anti-CD45-AF700 (HI30; BioLegend), anti-CD3-APC (SK7; BioLegend), anti-CD16-BV786 (3G8; Becton Dickinson), anti-CD103-BV650 (Ber-ACT8; Becton Dickinson), anti-CD4-BV605 (RPA-T4; Becton Dickinson), and anti-CD14-V500 (MφP9; Becton Dickinson) antibodies mixed with Staining Buffer (PBS 1X 3% FBS) containing 50% Brilliant Stain Buffer (Becton Dickinson) for 20 minutes at RT. Cells were then washed with Staining Buffer (SB) and resuspended with PBS 1X 2% BSA at a maximum concentration of 20 million/ml. Different trNK subsets were immediately sorted using the Cytek Aurora Cell Sorter. The purity of the cells was >95% in all cases. CD4^+^CD69^+^ T cells were also sorted and infected by spinoculation. Uninfected cells were also spinoculated with only R10 and served as an infection control. Sorted trNK cells were cocultured with autologous HIV-infected CD4^+^CD69^+^ T cells at effector-to-target (E:T) ratios of 1:1 to 1:3 in R20 medium supplemented with 100 U/ml IL-2. For suspension cocultures, cells were plated in 96-well round-bottom plates. In 2D and 3D collagen conditions, 96-well flat-bottom plates were coated with 10 µg/cm^2^ collagen I and incubated for 30 minutes to 2 hours at 37 °C in 5% CO_2_. For 3D collagen I gels (1.5 mg/ml), 15 µl of R20 medium with 100 U/ml IL-2 was mixed with 15 µl of neutralized collagen I (as described in the ‘*Extracellular matrix effects on the natural cytotoxicity of tonsil-resident NK cells*’ section). After collagen solidification (30 minutes to 2 hours at 37 °C, 5% CO_2_), 170 µl of R20 medium with IL-2 (100 U/ml) was added on top, and the medium was refreshed every 2-3 days. After 5 days of coculture, collagen-embedded cells were harvested using 3.3 mg/ml Collagenase IV (Gibco) and incubated for 30 minutes at 37 °C in 5% CO^2^. Subsequently, cells were stained for flow cytometry to assess the viability of remaining HIV-infected cells. Cells were stained with LIVE/DEAD AQUA viability (ThermoFisher) for 20 minutes at RT. After washing once with PBS 1X, cells were stained with anti-CD45-BV605 (HI30; Becton Dickinson), anti-CD3-AF700 (SK7; BioLegend), and anti-CD8-APC (RPA-T8; Becton Dickinson) antibodies for 20 minutes at RT. Alternatively, cells were stained with anti-CD45-AF700 (HI30; BioLegend), anti-CD3-APC (SK7; BioLegend), anti-CD8-BV650 (RPA-T8; Becton Dickinson), and anti-CD4-BV605 (RPA-T4; Becton Dickinson) antibodies for 20 minutes at RT. Cells were then washed with SB, fixed and permeabilized with Fixation/Permeabilization Solution (Becton Dickinson) for 20 minutes at 4°C, washed with BD Perm/Wash buffer and stained with anti-p24-PE (KC57; Beckman Coulter) for 30 min on ice followed by 30 min at RT. After washing, cells were fixed with PFA (2%) and acquired on a BD LSR Fortessa flow cytometer. Data were analyzed using FlowJo V10.

### Gene expression analysis by single-cell RNA-seq

Tonsillar cell suspensions were obtained from HIV-infected and uninfected tonsillar explants cultured for 5 days. CD56+ cells were enriched using the MagniSort™ Human CD56 Positive Selection Kit. After enrichment, the cells were washed and resuspended in 1X PBS for viability assessment with LIVE/DEAD Viability dye, incubated for 20 minutes. Subsequently, the cells were washed, resuspended in SB, and labeled with surface antibodies targeting lineage markers: anti-CD45-AF700 (HI30; BioLegend), anti-CD14-V500 (MφP9; Becton Dickinson), anti-CD19-APC-Fire750 (HIB19; BioLegend), anti-CD3-APC (SK7; BioLegend), anti-CD56-FITC (B159; Becton Dickinson). Finally, the stained cells were washed again in SB and resuspended in PBS containing 0.04% BSA. The CD14^−^CD19^−^CD3^−^CD56^+^ cell population was sorted from the tonsillar CD56+ cell suspensions using a FACS instrument (Cytek Aurora CS), achieving a final purity of >95% and 11,700 and 16,000 cells from HIV-infected and uninfected tissue blocks, respectively. The cellular concentration and viability were verified staining the sample with Trypan Blue (Fisher SC) and evaluated by microscopy. Individual cells were partitioned into Gel Bead⍰In⍰Emulsions by using the Chromium X System (10X Genomics). Single cell suspensions were converted to barcoded scRNA-seq libraries using the Chromium GEM-X Single Cell 3’ Reagent Kits v4 (10X Genomics) following manufacturer’s instructions. After cell lysis and barcoded reverse transcription of RNA, cDNA was amplified during 12 cycles and cDNA QC and quantification were performed on an Agilent TapeStation D5000 DNA ScreenTape (Agilent Technologies). cDNA libraries were indexed by PCR during 15 cycles using PN-3000431 Chromium Dual Index Kit TT Set A. Size distribution and concentration of 3’ cDNA libraries were verified on an Agilent TapeStation D5000 DNA ScreenTape (Agilent Technologies). Finally, sequencing cDNA libraries was carried out on a Nova Seq 6000 sequencer (Illumina) to obtain approximately 40,000 paired-end 150 bp reads per cell. The Cell Ranger pipeline (v8.0.1) was used for alignment, filtering, barcode counting, and UMI counting. Quality controls were conducted with FastQC v0.11.7 to ensure suitability for downstream analysis. Cells were filtered based on >200 detected features, <20% mitochondrial counts, and >5% ribosomal reads. Potential doublets (>50,000 UMI counts) were excluded. Sequencing reads were mapped to the human reference genome, and gene expression was quantified using the Cell Ranger count tool. Data normalization was performed using the SCTransform method. Principal Component Analysis (PCA) on the top 3,000 variable features was used for dimensionality reduction, retaining the first 30 principal components. Clustering was performed with an SNN modularity optimization algorithm, and cell distribution was visualized via UMAP. Integration of samples was achieved with the Harmony algorithm. Cluster marker identification and differential expression analyses were conducted using the FindMarkers function in the Seurat package, with a resolution of 0.7. Automatic cell type annotation using the SingleR method (v1.10) with MonacoImmuneData as reference identified a small cluster (1.5% of total CD3^−^CD56^+^ cells) as B cells and it was excluded from subsequent analysis, consistent with previous scRNA-seq studies of tonsillar ILCs that encountered similar contamination issues^70^. Biological significance was assessed via enrichment analysis using the Gene Ontology (GO) Biological Process category and Reactome Pathway Knowledgebase. Differentiation trajectory analysis was performed with the Slingshot package v2.4. Bioinformatics analyses were conducted at the Statistics and Bioinformatics Unit (UEB) at Vall d’Hebron Research Institute (VHIR).

### Dimensionality Reduction Analysis

Dimensionality reduction analysis of flow cytometry data was performed using the OMIQ software from Dotmatics (https://www.omiq.ai/, www.dotmatics.com). All events within pre-gated Live CD45^+^CD14^−^CD19^−^CD3^−^CD56^+^ NK cells were concatenated for each group per culture condition and analyzed. For NK cell phenotyping, cell clusters were identified by Flow-Self Organizing Maps (FlowSOM) artificial intelligence algorithm based on the expression of CD16, CD49a, CD69, CD103, KIRs, and NKG2C. Next, optimized T-distributed stochastic neighbor embedding (opt-SNE) analysis was employed to depict the identified clusters. The frequency of marker expression in each cluster was represented in a heatmap generated using GraphPad Prism software.

### Statistical Analysis

Statistical tests were performed with GraphPad Prism software (version 8.3) and are reported within each figure caption. Skillings-Mack test was performed on XLSTAT software. *P* values <0.05 were considered statistically significant. A minimum threshold of 10-15 cells was set to ensure robust measurement of marker expression by flow cytometry.

### Materials availability

This study did not generate new unique reagents. All reagents used are indicated in this section and are available.

### Data availability

All data associated with this study are provided within the paper or the supplemental information, and raw data are included in the Supporting Data Values file. Sequencing data are available in the NCBI GEO under accession number GSE286121. All data reported in this paper will be shared by the lead contact upon request. Due to the sensitivity of the data, individual participant data will not be shared.

This paper does not report original code.

Any additional information required to reanalyze the data reported in this paper is available from the lead contact upon request.

## AUTHOR CONTRIBUTIONS

M.B. designed, directed, and interpreted experiments. D.P. and A.G designed, performed, and interpreted experiments, analyzing the resulting data. N.S-G. set up the tonsillar explant model. F.P., N. O., I. L., J. L and V.F. were responsible for surgical resections, specimen handling and storage, and related clinical data collection. M.G interpreted experiments. D.P. and M.B. wrote the initial manuscript, and all of us contributed to the editing of the manuscript.

## ACKNOWLEDGMENTS

The project leading to these results has received funding from “la Caixa” Banking Foundation under the project code [LCF/PR/ HR20-00218]. This study was supported by the Agencia Estatal de Investigación project PID2021-123321OB-I00 funded by MCIN/AEI/10.13039/501100011033/FEDER, UE; The Spanish “Ministerio de Economía y Competitividad, Instituto de Salud Carlos III” (ISCIII, PI20/00160); and the Gilead fellowship GLD22/00152. M.B is supported by the Miguel Servet program funded by the Spanish Health Institute Carlos III (CPII22/00005). D.P was supported by the VHIR 2020 Ph.D. Fellowship. The funders had no role in study design, data collection, and analysis, the decision to publish, or preparation of the manuscript.

## RESOURCES TABLE

| REAGENT or RESOURCE | SOURCE | IDENTIFIER |
| --- | --- | --- |
| **Antibodies** |  |  |
| FITC anti-human CD56 Clone B159 | Becton Dickinson | Cat#562794; RRID:AB_2737799 |
| PE-Cy7 anti-human CD49a Clone TS2/7 | BioLegend | Cat#328312; RRID:AB_2566272 |
| PE-Cy5.5 anti-human CD158a, h Clone EB6B | Beckman Coulter | Cat#A66898; RRID:AB_2857330 |
| PE-Cy5.5 anti-human CD158b1/b2, j Clone GL183 | Beckman Coulter | Cat#A66900; RRID:AB_2857331 |
| PE-CF594 anti-human CD69 Clone FN50 | Becton Dickinson | Cat#562645; RRID:AB_2737699 |
| PE anti-human NKG2C Clone 134591 | R&D Systems | Cat#FAB138P; RRID:AB_2132983 |
| APC anti-human CD3 Clone SK7 | BioLegend | Cat#344812; RRID:AB_10645473 |
| BV786 anti-human CD16 Clone 3G8 | Becton Dickinson | Cat#563690; RRID:AB_2744299 |
| BV650 anti-human CD103 Clone Ber-ACT8 | Becton Dickinson | Cat#743653; RRID:AB_2741655 |
| BV605 anti-human CD45 Clone HI30 | Becton Dickinson | Cat#564047; RRID:AB_2744403 |
| V500 anti-human CD14 Clone MφP9 | Becton Dickinson | Cat#562693; RRID:AB_2737727 |
| V500 anti-human CD19 Clone HIB19 | Becton Dickinson | Cat#561121; RRID:AB_10562391 |
| APC-Fire750 anti-human CD19 Clone HIB19 | BioLegend | Cat#302258; RRID:AB_2629691 |
| BV421 anti-human CD107a Clone H4A3 | Becton Dickinson | Cat#562623; RRID:AB_2737685 |
| AF700 anti-human IFN-γ Clone B27 | ThermoFisher | Cat#MHCIFG29; RRID:AB_2539767 |
| BB700 anti-human c-kit Clone YB5.B8 | Becton Dickinson | Cat#566548; RRID:AB_2744268 |
| APC-Vio770 anti-human CRTH2 Clone REA598 | Miltenyi Biotec | Cat#130-119-623; RRID:AB_2751763 |
| AF700 anti-human CD45 Clone HI30 | BioLegend | Cat#304024; RRID:AB_493760 |
| BV605 anti-human CD127 Clone A019D5 | BioLegend | Cat#351334; RRID:AB_2562022 |
| BV421 anti-human NKp44 Clone p44-8 | Becton Dickinson | Cat#744299; RRID:AB_2742129 |
| AF700 anti-human CD3 Clone SK7 | BioLegend | Cat#344822; RRID:AB_2563420 |
| BV605 anti-human CD4 Clone RPA-T4 | Becton Dickinson | Cat#562658; RRID:AB_2744403 |
| BV650 anti-human CD8 Clone RPA-T8 | Becton Dickinson | Cat#563821; RRID:AB_2744462 |
| APC anti-human CD8 Clone RPA-T8 | Becton Dickinson | Cat#555369; RRID:AB_398595 |
| PE anti-HIV p24 Clone KC57-RD1 | Beckman Coulter | Cat#6604667; RRID:AB_1575989 |
| BB700 anti-human CXCR5 Clone RF8B2 | Becton Dickinson | Cat#566469; RRID:AB_2869769 |
| PerCP-Cy5.5 anti-human CD137 Clone 4B4-1 | BioLegend | Cat#309818; RRID:AB_2207741 |
| APC-Fire750 anti-human PD-1 Clone EH12.2H7 | BioLegend | Cat#329954; RRID:AB_2616721 |
| APC-Cy7 anti-human CD25 Clone M-A251 | Becton Dickinson | Cat#557753; RRID:AB_396859 |
| AF647 anti-human TIGIT Clone MBSA43 | ThermoFisher | Cat#51-9500-42; RRID:AB_2744746 |
| APC anti-human NKG2A Clone Z199 | Beckman Coulter | Cat#A60797; RRID:AB_10643105 |
| BV421 anti-human LAIR-1 Clone DX26 | Becton Dickinson | Cat#744557; RRID:AB_2742321 |
| BV421 anti-human KLRG1 Clone 14C2A07 | BioLegend | Cat#368604; RRID:AB_2566593 |
| **Virus strains** |  |  |
| Plasmid encoding HIV-1<sub>BaL</sub> | NIH AIDS Reagent Program | Cat#ARP-510 |
| **Chemicals, Peptides, Enzymes, and Recombinant Proteins** |  |  |
| Dulbecco's PBS (10X) w/o Ca & Mg, w/o Phenol Red | Capricorn Scientific | Cat#PBS-10XA |
| RPMI-1640 | Fisher Scientific | Cat#12004997 |
| Fetal Bovine Serum | Life Technologies | Cat#A5256701 |
| Bovine Serum Albumin | Merck | Cat#A7906-10G |
| Brilliant Stain Buffer | Becton Dickinson | Cat#566349 |
| BD GolgiPlug Protein Transport Inhibitor | Becton Dickinson | Cat#555029 |
| BD GolgiStop Protein Transport Inhibitor | Becton Dickinson | Cat#554724 |
| Paraformaldehyde 4% | ThermoFisher | Cat#J19943.K2 |
| NaOH 50% H<sub>2</sub>O | Sigma Aldrich | Cat#415413 |
| HEPES buffer solution 1M | Capricorn Scientific | Cat#HEP-B |
| Penicillin-Streptomycin | Capricorn Scientific | Cat#PS-B |
| Gentamicin | Gibco | Cat#15750037 |
| Amikacin | Vall d'Hebron Pharmacy |  |
| Amphotericin B | Gibco | Cat#15290026 |
| Timentin | Caisson Labs | Cat#11192002 |
| Human IL-2 | Vall d'Hebron Pharmacy |  |
| Collagenase IV | Fisher Scientific | Cat#10780004 |
| Rat tail Collagen I | Merck | Cat#08-115 |
| Collagen IV from Engelbreth-Holm-Swarm murine sarcoma basement membrane | Merck | Cat#C0543 |
| **Critical Commercial Kits** |  |  |
| LIVE/DEAD® Fixable Aqua Dead Cell Stain Kit | ThermoFisher | Cat#L34966 |
| AO/PI Cell Viability Kit | Isogen | Cat#F23001 |
| Britelite plus Reporter Gene Assay System | Revvity | Cat#6066761 |
| BD Cytotfix/Cytoperm™ | Becton Dickinson | Cat#554722 |
| EasySep™ Human CD19 Positive Selection Kit II | StemCell | Cat#17854 |
| EasySep™ Human CD56 Positive Selection Kit II | StemCell | Cat#17855 |
| CD56<sup>+</sup> Microbeads | Miltenyi Biotec | Cat#130-050-401 |
| Dynabeads™ CD4 Positive Isolation Kit | ThermoFisher | Cat#11145D |
| **Experimental Models: Cell Lines** |  |  |
| K562 human myelogenous leukemia cell line | Merck | Cat#89121407 |
| 293T cell line | ATCC | Cat#CRL-3216 |
| TZM-bl cell line | NIH AIDS Reagent Program | Cat#8129 |

| Software |  |  |
| --- | --- | --- |
| FlowJo | Becton Dickinson | https://www.flowjo.com/ |
| OMIQ | Dotmatics | https://www.omiq.ai/ |
| GraphPad | Dotmatics | https://www.graphpad.com/ |
| XLSTAT | Lumivero | https://www.xlstat.com/en/ |
| CellRanger | 10X Genomics | https://www.10xgenomics.com/ |

---

**Supplementary Figure S1. Tissue-resident memory phenotype of tonsillar NK cells.** Expression of resident (CD69, CD49a, and CD103) and memory (KIRs and NKG2C) markers measured by flow cytometry in different tonsillar tissue specimens. **(A)** Gating strategy of tissue-resident memory NK cells in tonsils. **(B)** Fluorescent Minus One (FMO) controls of tissue-resident memory markers expressed on tonsillar NK cells. **(C)** Coexpression frequencies of memory markers within resident CD16^+^ and CD16^−^ NK subpopulations. Median with interquartile range is represented. Statistical comparisons were performed using the Skillings-Mack test. Significant comparisons *p*<0.05. Green circles: CD16^+^; Blue squares: CD16^−^.

**Supplementary Figure S2. Functionality of tonsil-resident memory NK cells.** Expression of the degranulation marker CD107a and intracellular IFN-γ measured by flow cytometry in different tonsillar tissue specimens in the steady state and upon stimulation with K562 cells. **(A)** Cytotoxic degranulation (% of CD107a^+^IFN-γ^−^ cells) by resident (left) CD16^+^ and (right) CD16^−^ NK cells. **(B)** Cytokine production (% of CD107a^+^IFN-γ^+^ cells) by resident (left) CD16^+^ and (right) CD16^−^ NK cells. **(C)** Cytotoxic degranulation (% of CD107a^+^IFN-γ^−^ cells) by (left) CD16^+^CD69^+^CD49a^+^ and (right) CD16^+^CD69^+^CD49a^−^ NK cells. **(D)** Cytokine production (% of CD107a^+^IFN-γ^+^ cells) by (left) CD16^−^CD69^+^CD49a^+^ and (right) CD16^−^CD69^+^CD49a^−^ NK cells. **(E)** Cytokine production (% of CD107a^+^IFN-γ^+^ cells) by (left) CD16^+^CD69^+^CD49a^+^ and (right) CD16^+^CD69^+^CD49a^−^ NK cells. **(F)** Cytokine production (% of CD107a^+^IFN-γ^+^ cells) by CD16^−^CD69^+^CD49a^−^ NK cells. Statistical comparisons were performed using the Wilcoxon matched-pairs signed rank test and Skillings-Mack test when appropriate. Significant comparisons *p*<0.05. Green circles: CD16^+^; Blue squares: CD16^−^.

**Supplementary Figure S3. Immune checkpoint expression patterns in tonsil-resident NK cells.** Expression levels of the inhibitory immune checkpoints NKG2A, LAIR-1, TIGIT, PD-1, and KLRG1 were analyzed by flow cytometry in tonsillar tissue samples. **(A)** Flow cytometry examples of immune checkpoint expression within NK cells. **(B)** Frequencies of immune checkpoint expression within resident memory CD16^+^ NK cells. **(C)** Frequencies of immune checkpoint expression within resident memory CD16^−^ NK cells. Statistical comparisons were performed using the Friedman’s test or the Skillings-Mack test, as appropriate. Significant comparisons *p*<0.05. Green circles: CD16^+^; Blue squares: CD16^−^.

**Supplementary Figure S4. Correlations of tonsil-resident NK subpopulations with HIV infection.** Tonsillar tissue explants were cultured on hemostatic sponges and *ex vivo*-infected with HIV_Bal_ for 5 days. The expression of resident memory markers and immune checkpoints was measured by flow cytometry. **(A)** HIV *ex vivo* infection levels measured as the % of p24^+^ cells within CD3^+^CD8^−^ T cells. **(B-E)** Spearman correlations and correlation matrices showing the relationship between HIV p24 levels in explants and the expression of LAIR-1 on **(B-C)** CD16^+^ and **(D-E)** CD16^−^ tonsil-resident memory NK cells on day 0. Significant Spearman’s *p*<0.05. Green circles: CD16^+^; Blue squares: CD16^−^.

**Supplementary Figure S5. Gene expression profiles distinguish ILC clusters in tonsillar explants.** Tonsillar tissue explants were cultured on hemostatic sponges and *ex vivo*-infected with HIV_Bal_ for 5 days. The CD14^−^CD19^−^CD3^−^CD56^+^ cell population from HIV-uninfected and HIV-infected explants was isolated by FACS and analyzed using scRNA-seq. Heatmap showing the top 10 most upregulated genes in each ILC cell cluster.

**Supplementary Figure S6. Gene expression of canonical NK cell markers across ILC clusters.** Tonsillar tissue explants were cultured on hemostatic sponges and *ex vivo*-infected with HIV_Bal_ for 5 days. The CD14^−^CD19^−^CD3^−^CD56^+^ cell population from HIV-uninfected and HIV-infected explants was isolated by FACS and analyzed using scRNA-seq. **(A-G)** Violin plots illustrating the expression distribution of canonical human NK cell markers across identified clusters. Markers are grouped into the following categories: **(A)** NK-ILCs, **(B)** NK receptors, **(C)** resident markers, **(D)** ECM remodeling and NK invasion, **(E)** effector mediators, **(F)** chemokine receptors, and **(G)** cytokine receptors. The maximum gene expression value displayed is 7.

**Supplementary Figure S7. Distinct transcriptional signatures in tonsil-resident ILC/NK cells during early HIV exposure.** Tonsillar tissue explants were cultured on hemostatic sponges and *ex vivo*-infected by HIV_Bal_ for 5-6 days. The CD14^−^CD19^−^CD3^−^CD56^+^ cell population from HIV-uninfected and HIV-infected explants was isolated by FACS and analyzed using scRNA-seq. **(A)** Volcano plots of clusters 1, 3, 5, 6, 7, 8, 9, and 10 showing DEGs between uninfected and infected tissue explant CD56^+^ cells. For clusters 9 and 10, note that the volcano plots were generated using unadjusted *p*-values due to their low frequency representation. **(B)** Gene Ontology (GO) and Reactome pathway analyses highlighting selected pathways enriched in clusters 1, 3, 5, 6, 7, 8, 9, and 10 in response to HIV exposure. DEGs represented in each enriched pathway are shown.

**Supplementary Figure S8. Widespread enrichment of the long non-coding RNA *ENSG00000289901* across ILC clusters following HIV exposure.**

Tonsillar tissue explants were cultured on hemostatic sponges and *ex vivo*-infected with HIV_Bal_ for 5 days. The CD14^−^CD19^−^CD3^−^CD56^+^ cell population from HIV-uninfected and HIV-infected explants was isolated by FACS and analyzed using scRNA-seq. **(A)** Violin plots illustrating *ENSG00000289901* expression across clusters 0 to 9 following HIV exposure. Corresponding log_2_ fold-change values and adjusted *p*-values are provided in **Supplementary Table S2**. Light color represents gene expression in HIV-uninfected samples, while bold color indicates gene expression in HIV-infected samples. **(B)** Predicted functional interactions of *ENSG00000289901* with proteins, displaying the top 10 interactions identified using CATrapid omics v2.1.

**Supplementary Figure S9. Early HIV exposure of tissue explants does not impact on the frequency distribution of tonsil-resident NK subsets.** Tonsillar tissue explants were cultured on hemostatic sponges and *ex vivo*-infected with HIV_Bal_ for 5 days. The expression of resident and memory markers was measured by flow cytometry in the steady state and upon stimulation with K562 cell coculture from uninfected and infected tissue explant cells. **(A-D)** Frequencies of **(A)** total CD56^+^ NK cells, **(B)** CD16^+/−^ NK cell subsets, **(C)** resident CD16^+/−^ NK cells, and **(D)** memory CD16^+/−^ NK cells in uninfected and infected tissue explants. Statistical comparisons were performed using the Wilcoxon matched-pairs signed rank test. Green circles: CD16^+^; Blue squares: CD16^−^.

**Supplementary Table S1. Donor characteristics of human lymphoid tissue samples.**

Sample, age and sex are provided.

**Supplementary Table S2. Differentially expressed genes in each cluster following 5 days of HIV exposure in tonsillar explants.**

FC, log_2_ fold-change of the gene expression in HIV-exposed cells vs HIV-unexposed per cluster. Adjusted *p*-value is based on Bonferroni correction and shown as −log_10_.

