## Supplementary Figures and Tables for "Early resident NK cell response to local HIV infection in lymphoid tissue"

#### Supplementary Figure S1

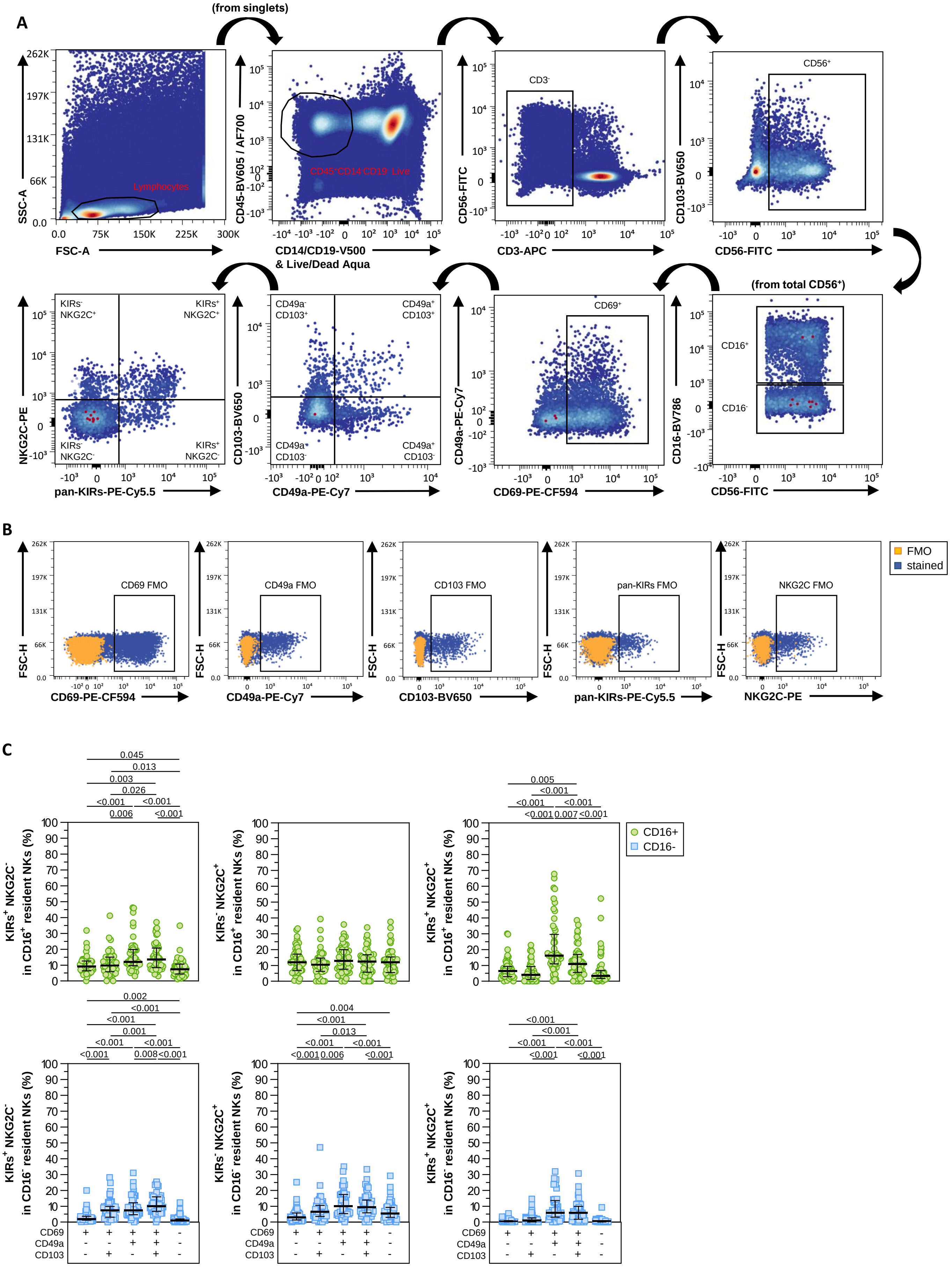

#### Supplementary Figure S2

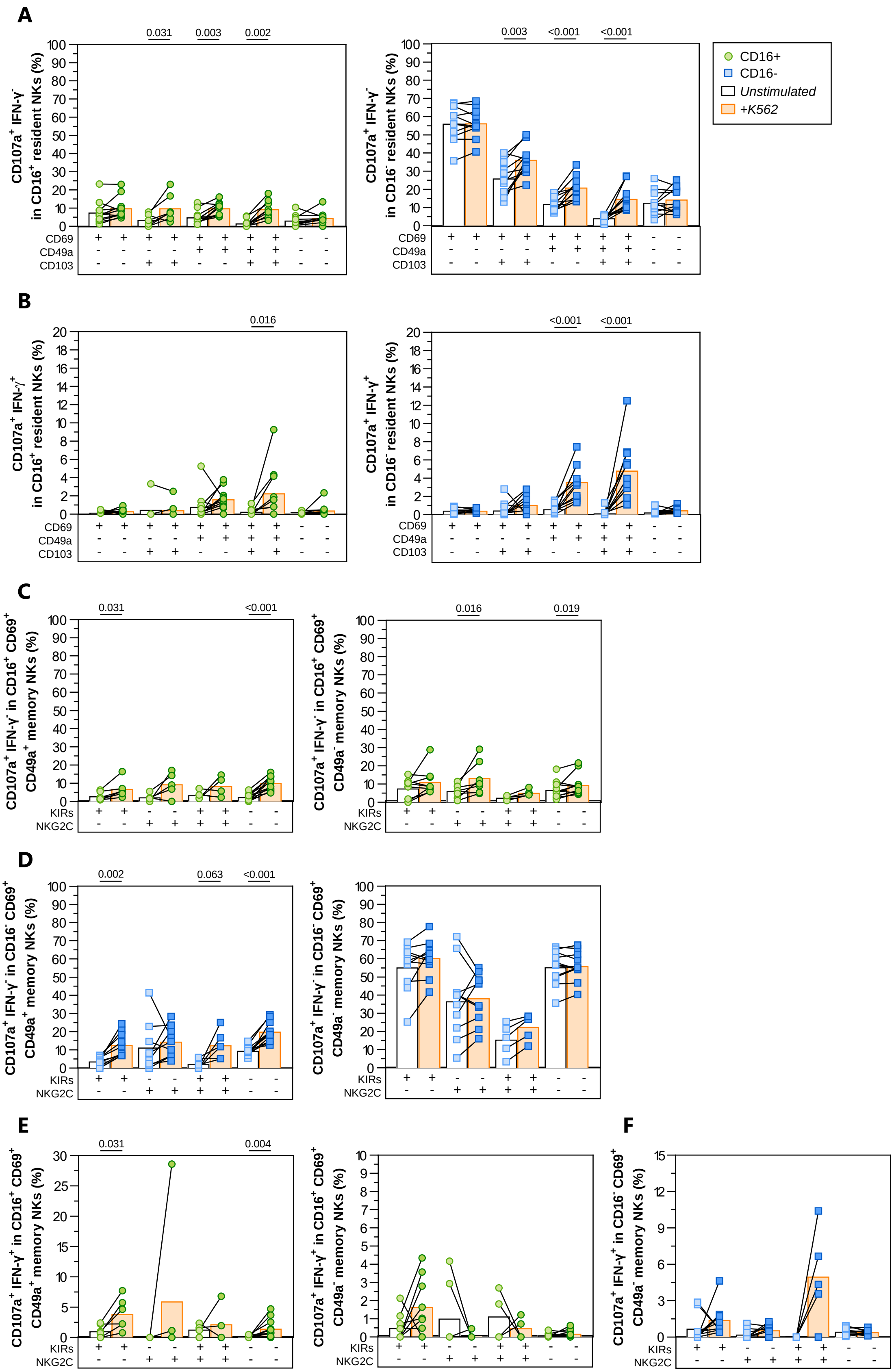

#### Supplementary Figure S3

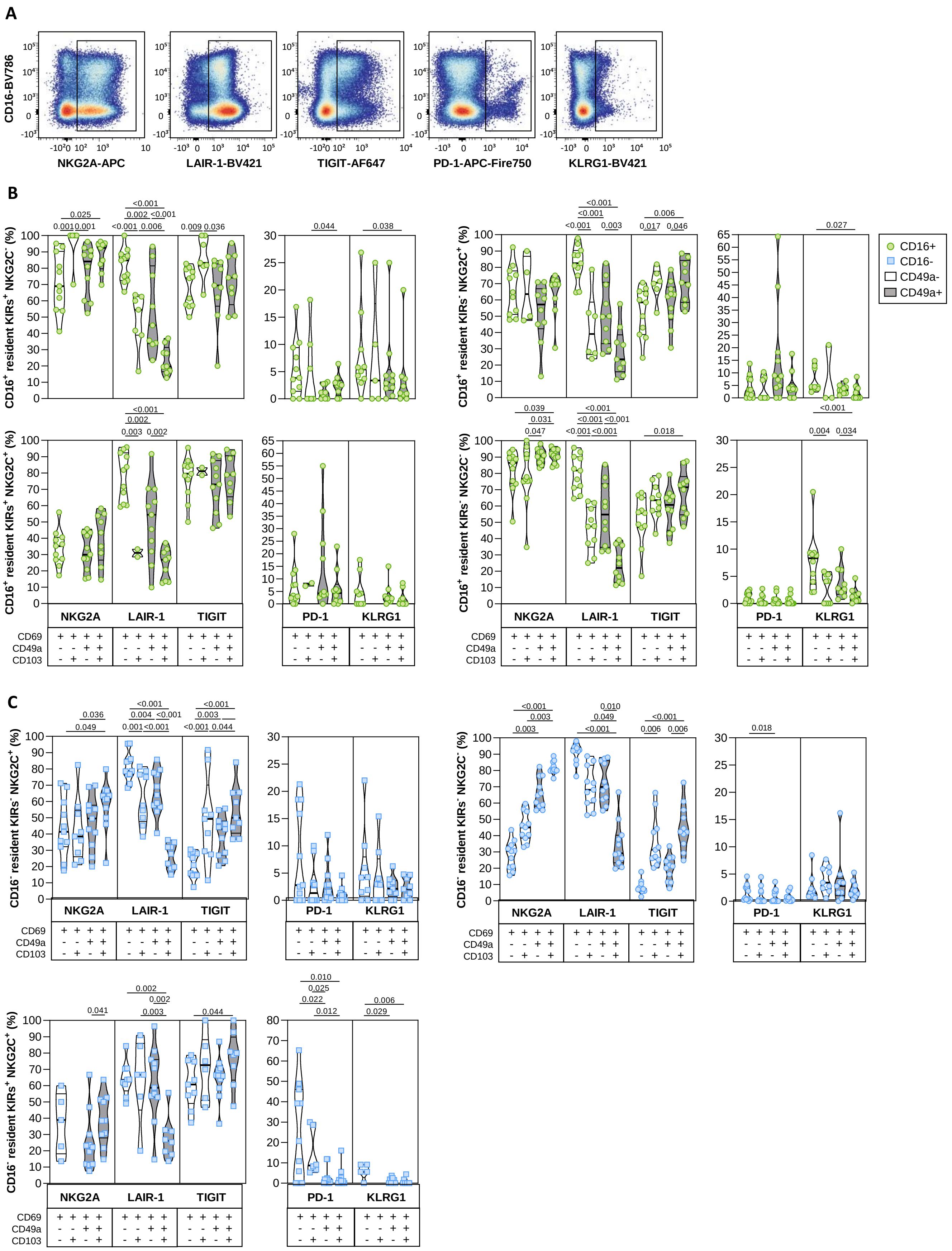

### Supplementary Figure S4

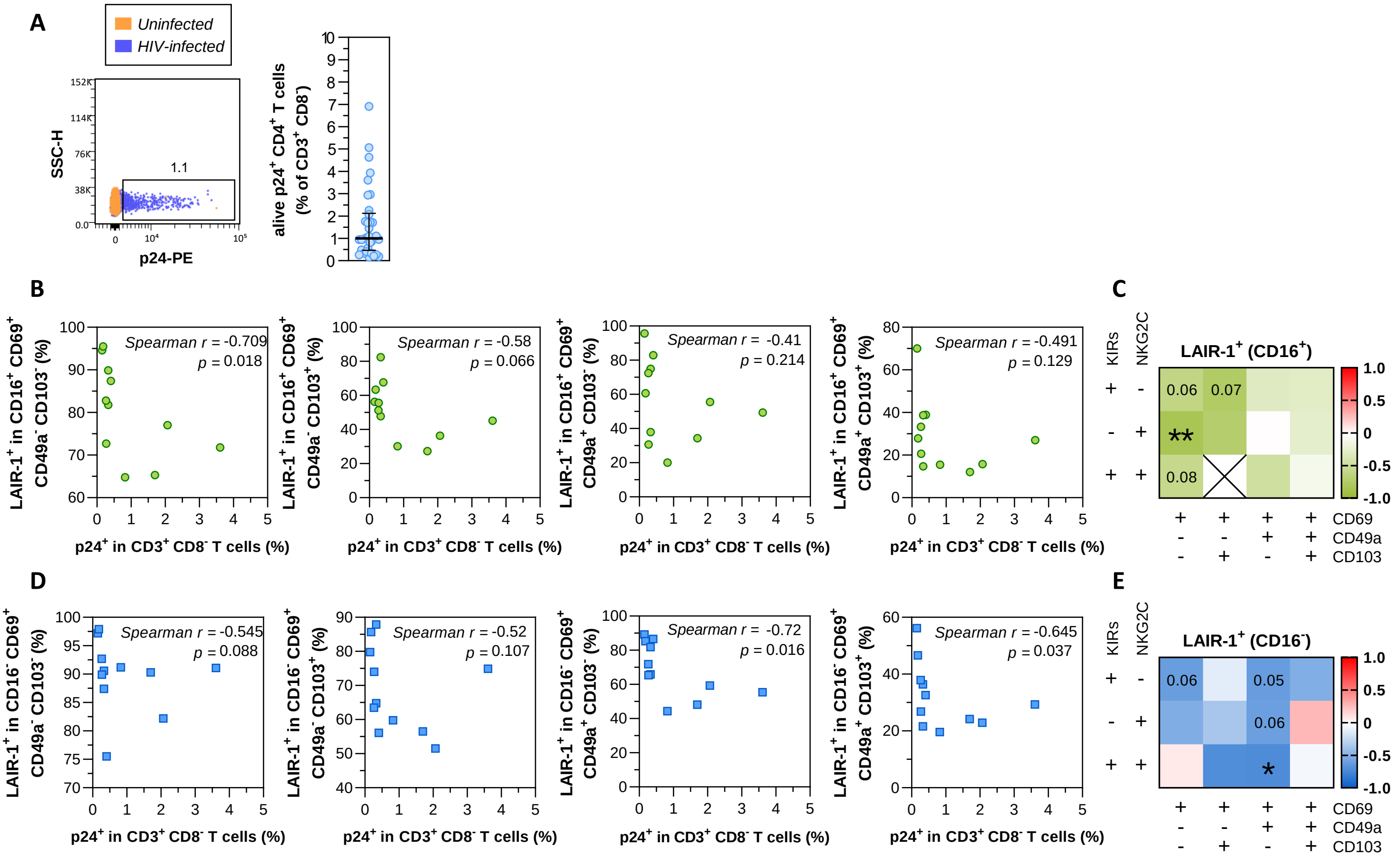

**Supplementary Figure S5**

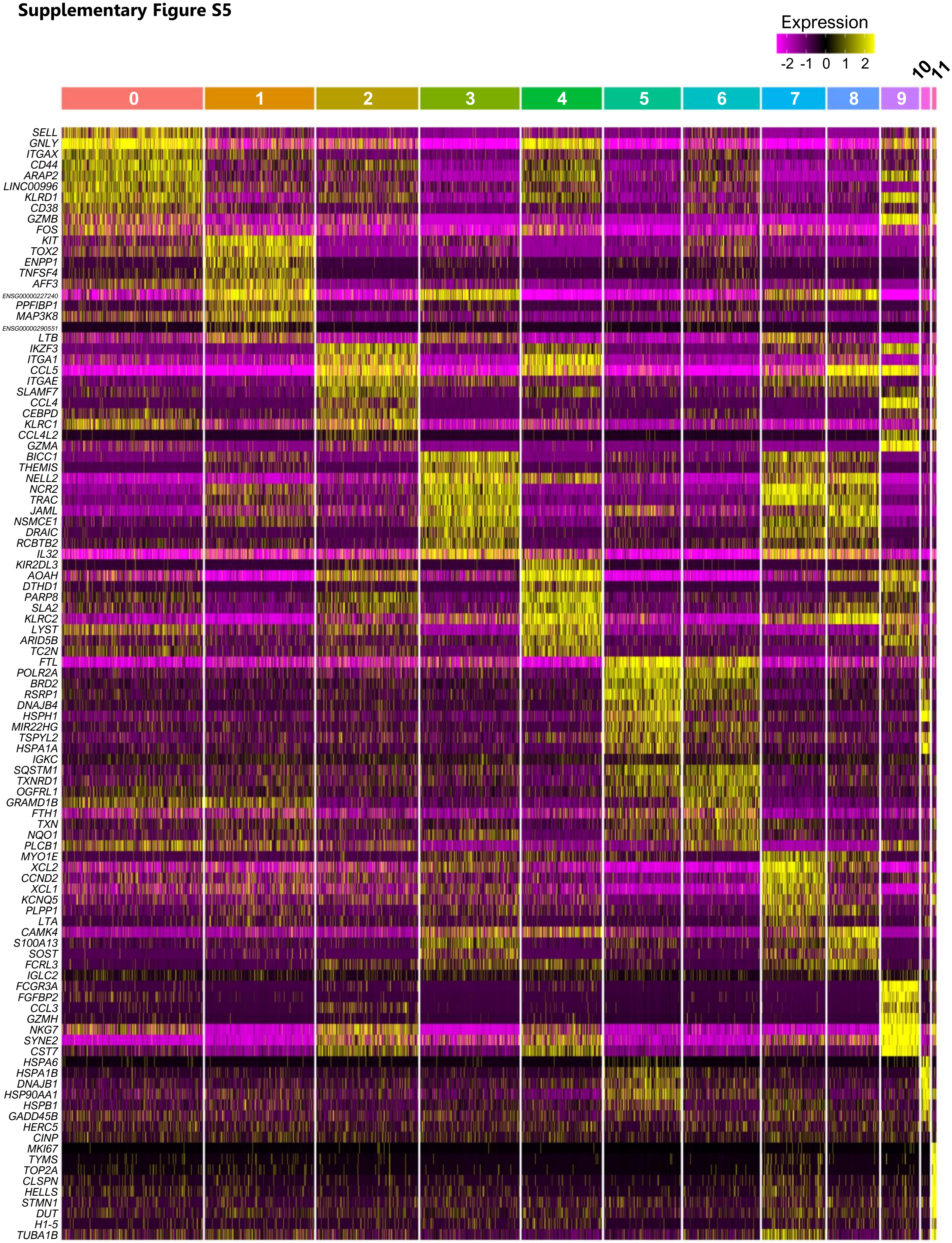

#### Supplementary Figure S6

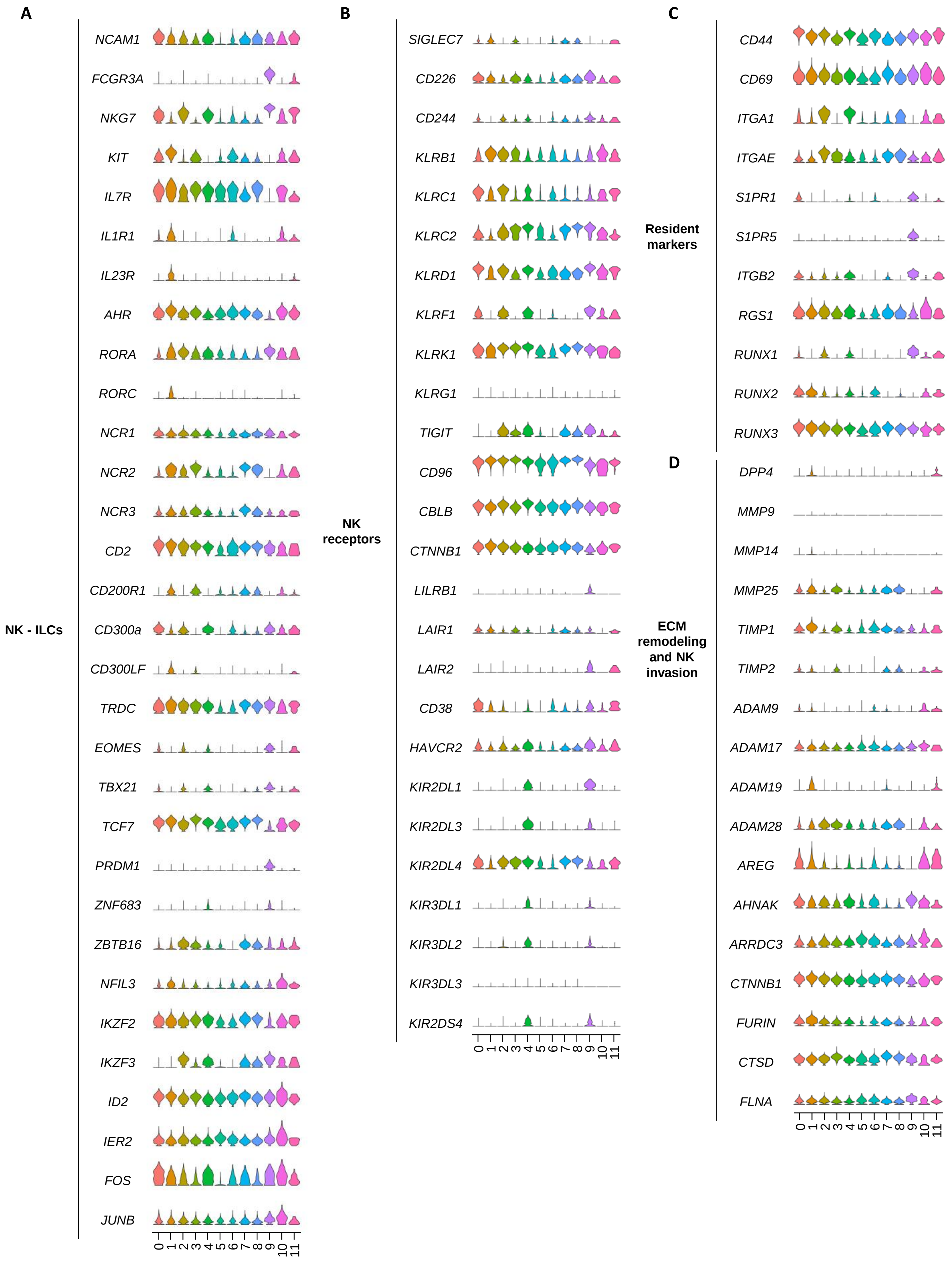

Supplementary Figure S6 *(continued)*

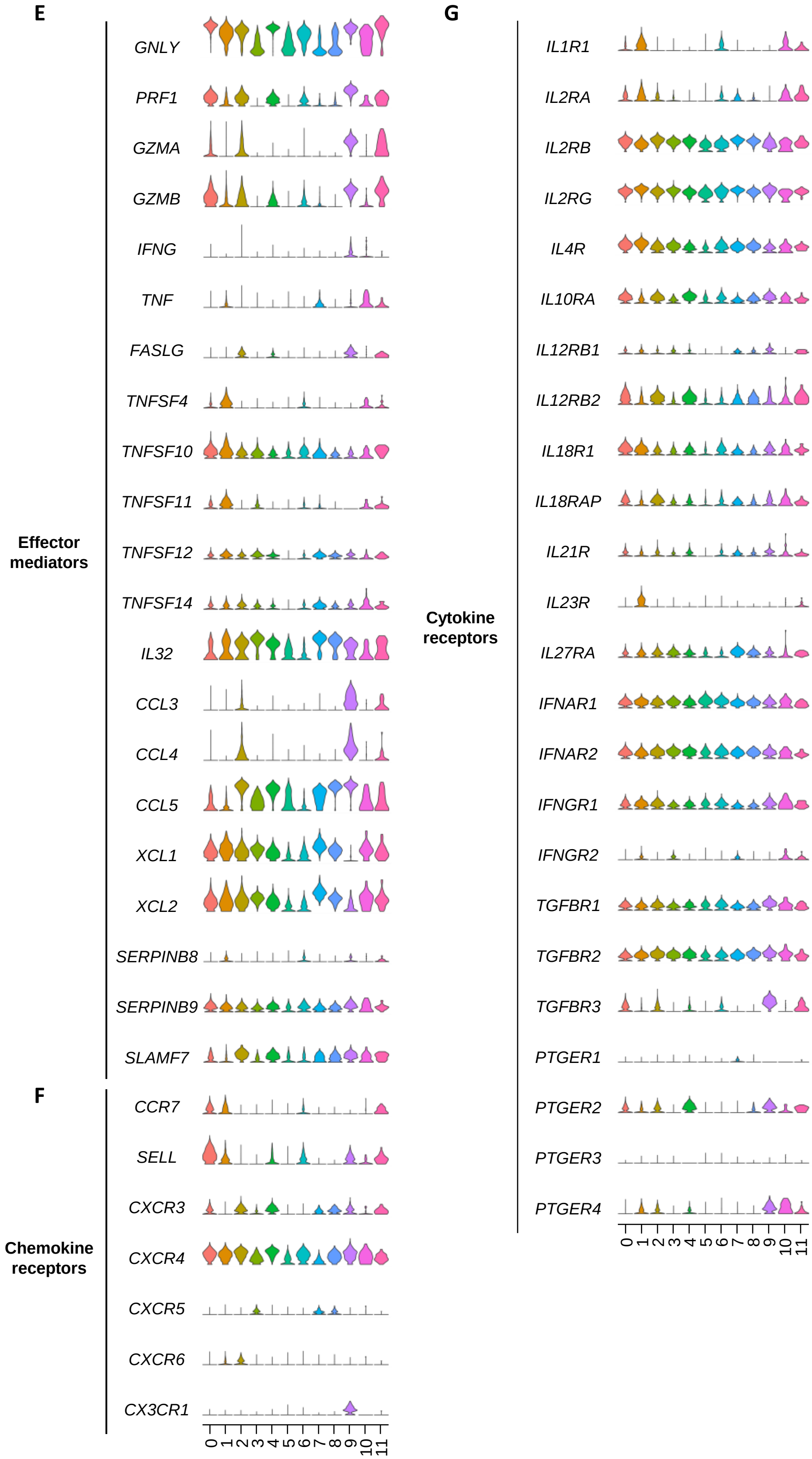

#### Supplementary Figure S7

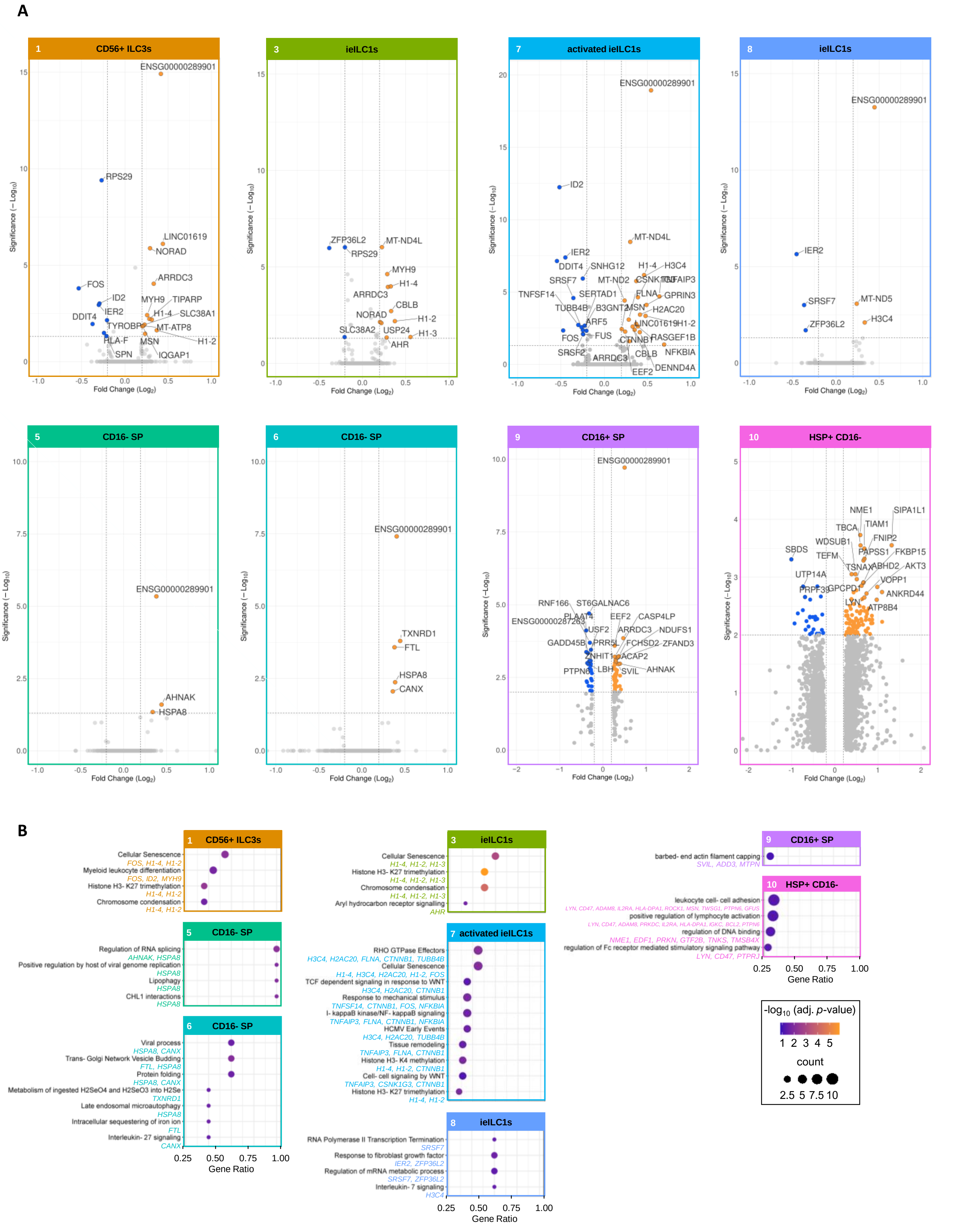

### Supplementary Figure S8

**A**

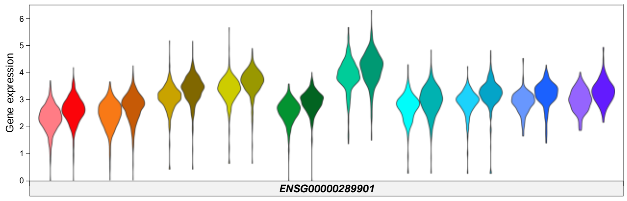


**B**

| Top 10 RNA-binding domains for lncRNA *ENSG00000289901* | | |
|---|---|---|
| **Protein** | **Interaction Propensity** | **Z score** |
| SEC63 | 85.04 | 5.31 |
| TCERG1 | 83.22 | 5.18 |
| SLC4A1AP | 81.91 | 5.09 |
| ZFC3H1 | 81.88 | 5.09 |
| ESF1 | 81.88 | 5.09 |
| L1TD1 | 81.71 | 5.08 |
| ATXN1 | 81.34 | 5.05 |
| BAZ2B | 81.09 | 5.03 |
| HTATSF1 | 80.55 | 5.00 |
| SUZ12 | 79.70 | 4.94 |

#### Supplementary Figure S9

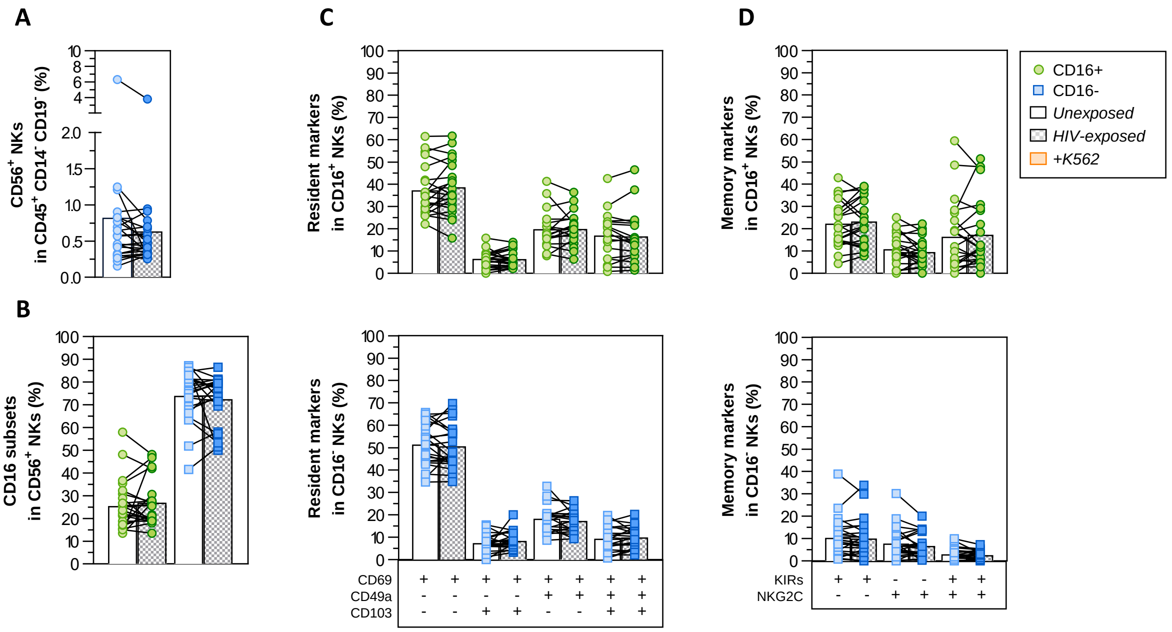

**Supplementary Table S1.** Donor characteristics of human lymphoid tissue samples.

| Sample (%) | Age mean (IQR) | Sex (%) |
|---|---|---|
| **Tonsil (93.06)**<br>**Adenoid (6.94)** | 5.98 (3.10) | Female (31.94)<br>Male (68.06) |

IQR: Interquartile range

**Supplementary Table S2.** Differentially expressed genes in each cluster following 5 days of HIV exposure in tonsillar explants.

**A**

| Upregulated genes | | | |
|---|---|---|---|
| **Gene** | **Cluster** | **FC** | **Adjusted *p*-value (-log$_{10}$)** |
| *AHNAK* | 0 | 2.1 | 13.8 |
| | 5 | 2.8 | 25 |
| *AHR* | 3 | 1.9 | 12 |
| *ARRDC3* | 0 | 2.8 | 8.8 |
| | 1 | 2.2 | 8.5 |
| | 2 | 2 | 8.2 |
| | 3 | 2 | 7.3 |
| | 7 | 2 | 7.1 |
| *CANX* | 6 | 2.3 | 7 |
| *CBLB* | 3 | 2.2 | 5.1 |
| | 7 | 2.3 | 5.8 |
| *CD96* | 4 | 1.8 | 3.7 |
| *CSNK1G3* | 7 | 2.5 | 3 |
| *CTNNB1* | 7 | 2.1 | 3 |
| *DENND4A* | 7 | 2.6 | 9.4 |
| *ENSG00000289901* | 0 | 2.7 | 1.6 |
| | 1 | 2.6 | 22.9 |
| | 2 | 3 | 2.2 |
| | 3 | 2.6 | 10.2 |
| | 4 | 3.1 | 2.2 |
| | 5 | 2.4 | 3.9 |
| | 6 | 2.6 | 7.6 |
| | 7 | 3.5 | 2.9 |
| | 8 | 2.8 | 2 |
| | 9 | 3.2 | 2.4 |
| *FLNA* | 7 | 1.9 | 3.1 |
| *FTL* | 6 | 2.4 | 1.4 |
| *GPRIN3* | 7 | 3.1 | 4 |
| *H1-2* | 1 | 2.3 | 1.4 |
| | 3 | 2.4 | 2.2 |
| | 7 | 3 | 2.7 |
| *H1-3* | 3 | 3.6 | 1.3 |
| *H1-4* | 1 | 2.1 | 10.1 |
| | 3 | 2.1 | 10.8 |
| | 7 | 2.9 | 24.4 |
| *H2AC20* | 7 | 2.6 | 4.9 |
| *H3C4* | 4 | 2.3 | 4.3 |
| | 7 | 2.4 | 4.9 |
| | 8 | 2.2 | 2.9 |
| *HSPA8* | 5 | 2.2 | 2.4 |
| | 6 | 2.4 | 2.6 |
| *LINC01619* | 1 | 2.8 | 2.1 |
| | 7 | 2.3 | 18.9 |
| *MT-ND4L* | 7 | 2 | 12.2 |
| *MYH9* | 0 | 1.9 | 7.4 |
| | 1 | 1.8 | 7.1 |
| | 3 | 1.9 | 8.5 |
| *NFKBIA* | 0 | 3.1 | 5.8 |
| | 2 | 2.6 | 4.7 |
| | 7 | 4.9 | 6.2 |
| *NORAD* | 0 | 1.9 | 4.6 |
| | 1 | 2 | 4.6 |
| *PLCG2* | 0 | 1.8 | 3.4 |
| | 2 | 1.9 | 4.1 |
| *RASGEF1B* | 7 | 2.5 | 3.3 |
| *SLC38A1* | 0 | 1.9 | 2.7 |
| *TIPARP* | 1 | 1.9 | 2.6 |
| *TNFAIP3* | 2 | 2.3 | 2.3 |
| | 7 | 4.4 | 2.4 |
| *TXNRD1* | 0 | 2.2 | 5.6 |
| | 6 | 2.8 | 13.3 |

**B**

| Downregulated genes | | | |
|---|---|---|---|
| **Gene** | **Cluster** | **FC** | **Adjusted *p*-value (-log$_{10}$)** |
| *DDIT4* | 0 | 3.3 | 1.8 |
| | 1 | 2.4 | 2.3 |
| | 2 | 3 | 2.1 |
| | 7 | 3.5 | 14.9 |
| *DUSP1* | 0 | 3.5 | 6.1 |
| | 4 | 3.7 | 5.9 |
| *DUSP2* | 0 | 3.3 | 4 |
| | 2 | 3.4 | 3.8 |
| | 4 | 4.3 | 3 |
| *FOS* | 0 | 3.9 | 2 |
| | 1 | 3.4 | 2.4 |
| | 4 | 5.1 | 1.9 |
| | 7 | 3 | 3 |
| *GADD45B* | 4 | 2 | 22.2 |
| *GIMAP7* | 0 | 2 | 6 |
| | 2 | 2.4 | 4.6 |
| | 4 | 2.4 | 4 |
| *ID2* | 1 | 2 | 2.1 |
| | 7 | 3.3 | 1.4 |
| *IER2* | 0 | 2.5 | 1.6 |
| | 1 | 2 | 5.3 |
| | 2 | 3.5 | 3.8 |
| | 4 | 4 | 3.6 |
| | 7 | 2.8 | 1.3 |
| | 8 | 2.8 | 7.4 |
| *JUNB* | 4 | 2 | 2.4 |
| *RPS29* | 0 | 1.8 | 3.1 |
| | 1 | 1.9 | 2.8 |
| *SRSF7* | 7 | 2.3 | 2.6 |
| | 8 | 2.3 | 2.6 |
| *TNFSF14* | 7 | 2 | 2.2 |
| *TUBB4B* | 2 | 1.9 | 1.4 |
| | 7 | 1.8 | 1.6 |
| *ZFP36L2* | 0 | 1.9 | 3 |
| | 3 | 2.4 | 5.3 |
| | 4 | 2.4 | 1.7 |
| | 8 | 2.2 | 2.1 |

FC, log$_2$ fold-change of the gene expression in HIV-exposed cells vs HIV-unexposed per cluster. Adjusted *p*-value is based on Bonferroni correction and shown as -log$_{10}$.
